# Layer and rhythm specificity for predictive routing

**DOI:** 10.1101/2020.01.27.921783

**Authors:** André M. Bastos, Mikael Lundqvist, Ayan S. Waite, Nancy Kopell, Earl K. Miller

## Abstract

In predictive coding, experience generates predictions that attenuate the feeding forward of predicted stimuli while passing forward unpredicted “errors”. Different models have different neural implementations of predictive coding. We recorded spikes and local field potentials from laminar electrodes in five cortical areas (V4, LIP, area 7A, FEF, and PFC) while monkeys performed a task that modulated visual stimulus predictability. Pre-stimulus predictions were associated with increased alpha/beta (8-30 Hz) power/coherence that fed back the cortical hierarchy primarily via deep-layer cortex. Unpredictable stimuli were associated with increases in spiking and in gamma-band (40-90 Hz) power/coherence that fed forward up the cortical hierarchy via superficial-layer cortex. Area 7A uniquely showed increases in high-beta (~22-28 Hz) power/coherence to unpredicted stimuli. These results suggest that predictive coding may be implemented via lower-frequency alpha/beta rhythms that “prepare” pathways processing predicted inputs by inhibiting feedforward gamma rhythms and associated spiking.

## Introduction

The brain exploits predictability. Predicted sensory inputs are not fully processed, making cortical processing more efficient and preventing “sensory overload”. Visuomotor integration, visual/auditory speech perception, and visual perception all benefit from predictability of sensory inputs (Arnal and Giraud, 2012; Bell et al., 2016; Körding and Wolpert, 2004). According to Predictive Coding theory (Friston, 2010; Mumford, 1992; Rao and Ballard, 1999), past sensory experience creates models that generate predictions that inhibit processing of expected inputs. Unexpected sensory inputs (“prediction errors”), being informative, are not inhibited and thus fully processed (and available to update the models). Indeed, all over visual cortex there is more spiking to unexpected stimuli (i.e., prediction errors, (Bell et al., 2016; Issa et al., 2018; Schwiedrzik and Freiwald, 2017; Zmarz and Keller, 2016) and higher fMRI BOLD signals (Alink et al., 2010; Kok et al., 2012; Summerfield et al., 2008)

Different models implement Predictive Coding differently, particularly in how predictions and prediction error signals are handled. Most current models propose separate circuits specialized for generating predictions and prediction errors (Bastos et al., 2012; Friston, 2010; Rao and Ballard, 1999; Spratling, 2008). Some propose that that prediction errors flow from higher to lower cortex only (Friston, 2010; Rao and Ballard, 1999) while others propose that prediction errors act locally within each cortical area or that predictions flow both up and down the cortical hierarchy (Spratling, 2008).

Another class of models is based on observations that alpha/beta (8-30 Hz) rhythms are more associated with top-down signaling and gamma (40-100 Hz) rhythms are more associated with bottom-up sensory inputs (Bastos et al., 2015; Buschman and Miller, 2007). The central idea is that alpha/beta feeds back predictions down the cortical hierarchy while gamma feeds forward unexpected inputs up the hierarchy. This is supported by human electro/magnetoencephalography (MEG/EEG) studies and a recent monkey electrocorticography study (Arnal and Giraud, 2012; Auksztulewicz and Friston, 2016; Bauer et al., 2006; Brodski et al., 2015; Chao et al., 2018; Mayer et al., 2016; van Pelt et al., 2016; Todorovic et al., 2011). Further, alpha/beta power is higher in deep-layer cortex and gamma power is higher in superficial layers (Bastos et al., 2018; Maier et al., 2010; Smith et al., 2013). This, plus recent evidence that deep-layer alpha/beta inhibits superficial layer gamma (Bastos et al., 2018) suggested a new implementation of predictive coding: Predictive Routing.

According to Predictive Routing, there are not separate circuits or specialized neurons for predictions vs prediction errors. Instead, predictions “prepare” the pathways in sensory cortex that process predicted sensory inputs. They do so when top-down alpha/beta rhythms in deep-layer cortex inhibit gamma rhythms/spiking in superficial layers that feedforward bottom-up inputs. Thus, there are no special error-detection circuits or mechanisms. There are instead pathways that are prepared/inhibited by alpha/beta prediction signals and thus can express gamma/spiking to unexpected inputs.

To test these models, we recorded LFPs and spiking using multi-area, multi-laminar recordings from a visual area (V4) and higher-order cortical areas (posterior parietal cortex and prefrontal cortex) simultaneously. We manipulated the predictability of objects used in a working memory task. This revealed layer and frequency-specific associations with predictions and prediction errors as well as evidence for the direction of flow of these signals. The results lend support for different aspects of previous predictive coding models as well as support for Predictive Routing.

## Results

### Task, behavior, and neurophysiological recordings

Monkeys performed a delayed match to sample (DMS) task (Figure 1A). The task was performed in one of two modes: unpredictable blocks-where one of 3 objects was chosen randomly as a sample on each trial for a block of 50 trials and predictable blocks - where the same object was used as a sample for 50 consecutive trials. The purpose of the DMS task was to ensure that animals were always engaged and attending to stimuli.

**Figure 1.**
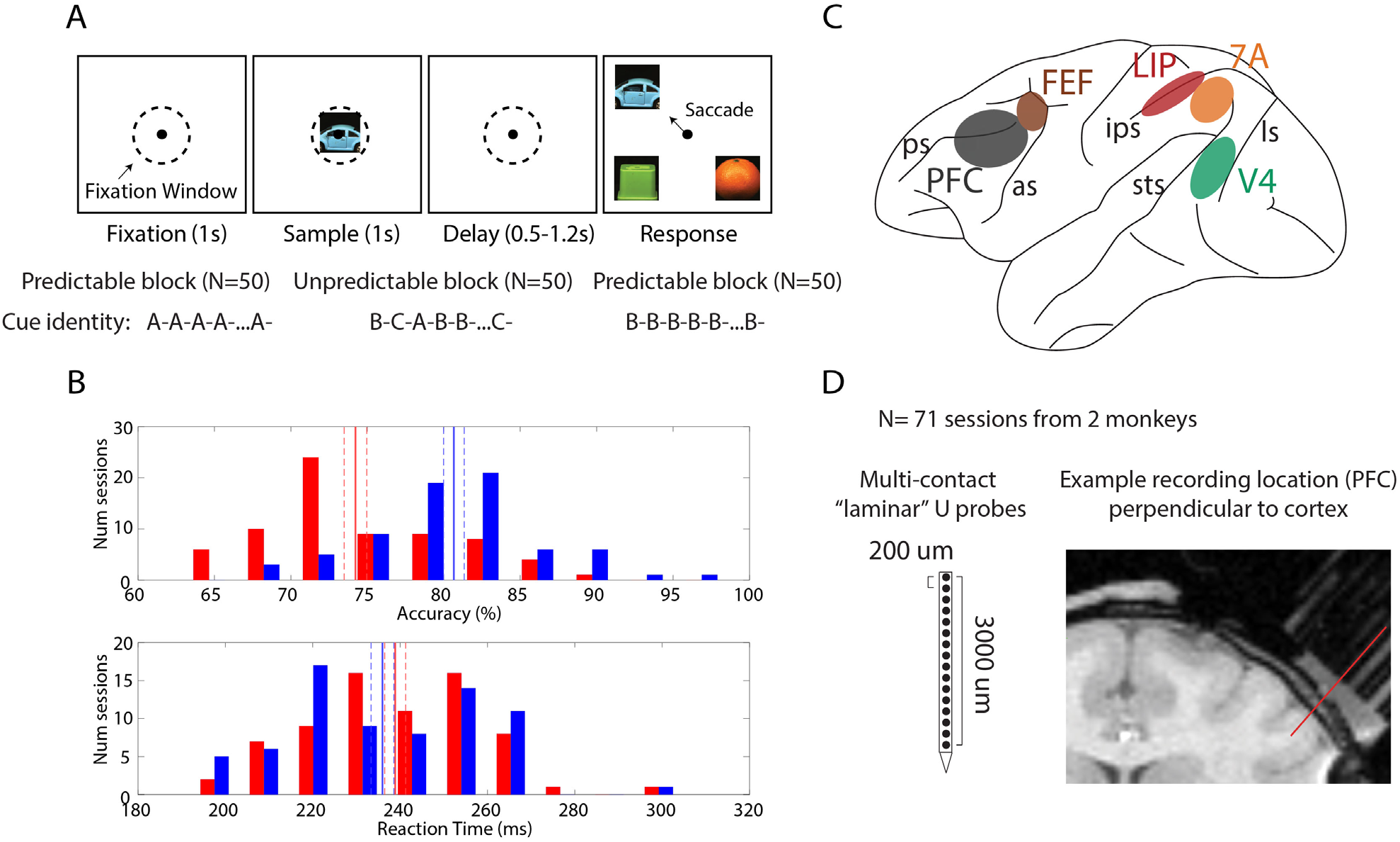
Task design, recordings, and behavior. A. Task design: after a one second fixation window, a sample stimulus (one of three pictures) is shown for one second. After a variable delay, the sample re-appears at one of 4 locations (always randomized). Monkeys saccade to the sampled stimulus. The sample identity was either randomized (unpredictable blocks) or held constant (predictable blocks). B. Behavioral performance across 65 sessions. Upper panel, accuracy on the task during predictable vs. unpredictable blocks. Lower panel, same, but for reaction time. Solid bars denote the mean performance +/- SEM across sessions. C. Recorded brain areas. PFC: Prefrontal Cortex, FEF: Frontal Eye Fields, LIP: Lateral Intraparietal Suluc, 7A: Posterior Parietal area 7A, V4: visual area 4. D. Multi-electrode 16 channel Plexon U/V probes with 200um site-to-site spacing. MRIs were used to select grid locations for laminar (perpendicular) access in areas located on cortical gyri: V4, 7A, and PFC. An example penetration in PFC is shown.

Choosing the match was more accurate and faster when the sample had been predictable. Figure 1B (upper panel) shows the distribution of average performance across sessions and monkeys for predictable vs. unpredictable cuing. Average performance during predictable blocks was 80.6% vs. 74.2% for unpredictable blocks (sign test across sessions, P<1E-8). Figure 1B (lower panel) shows the corresponding distribution for reaction time. Although the effect size was small, the match was found significantly more quickly during predictable vs unpredictable Blocks (mean RT predictable cuing: 236ms, mean RT unpredictable cuing: 239ms, non-parametric sign test across sessions, P=0.017).

We recorded spiking and local field potentials (LFPs) using multi-laminar electrodes (Figure 2D) in five cortical areas spanning sensory (V4), posterior parietal (Lateral Intraparietal area – LIP, and area 7A), and prefrontal cortex (PFC and Frontal Eye Fields – FEF, Figure 1C) in two monkeys over 71 sessions. For areas on cortical gyri (V4, 7A, and PFC), we introduced the electrodes perpendicular to cortex (Figure 1D) to resolve recordings into superficial (layer 2/3) vs. deep (layer 5/6). Data were aligned to the top of cortex, as this was the most robust metric that with minimal assumptions (see Methods). Areas FEF and LIP are located in sulci. Recordings there were not layer-resolved.

**Figure 2.**
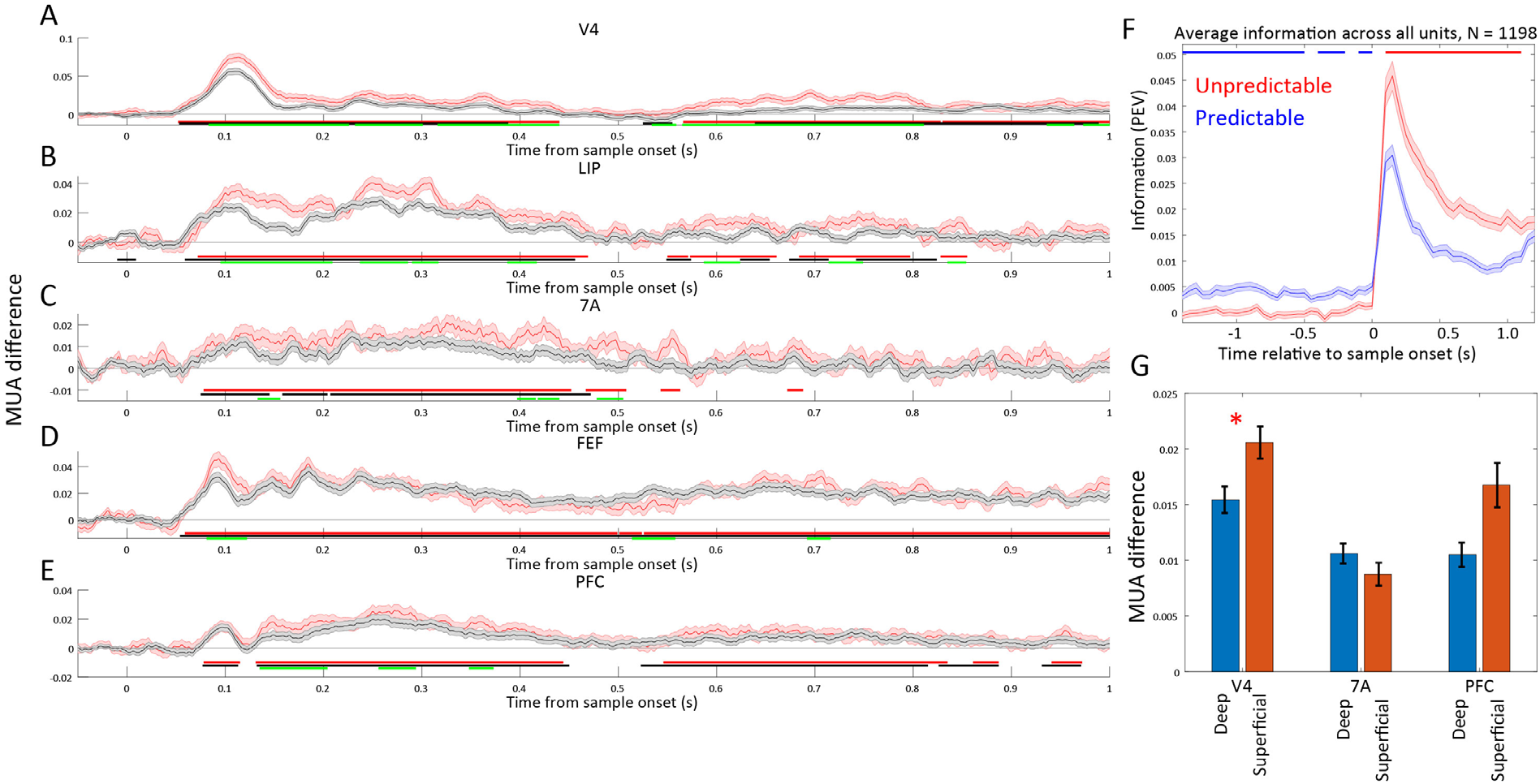
Spiking to unpredictable vs. predictable samples. A-E, Modulation of Multi-Unit Activity (MUA). Black lines: unpredictable (trials 11-50 in an unpredictable block) minus predictable (trials 11-50 of a predictable block) MUA relative to sample onset (time 0). Red lines: violation (trials 1-10 in an unpredictable block when it is preceded by a predictable block) minus predictable (trials 11-50 of a predictable block) MUA relative to sample onset (time 0). Mean across all available MUAs per area (N= 1,664, 736, 704, 880, 1472 for V4, LIP, 7A, FEF, PFC), +/- 1SEM across MUAs. Horizontal bars denote significance at p< 0.05 for unpredictable vs. predictable (black), violation vs. predictable (red), and violation vs. unpredictable (green). F. Mean information +/- SEM, quantified with Percent Explained Variance (PEV) in thresholded single units across all areas about the sample during predictable (blue line) vs. unpredictable (red line) trials. Horizontal blue bars indicate significant (p<0.05) unpredictable < predictable information. Horizontal red bars indicate significant (p<0.05) unpredictable > predictable information. G. Unpredictable minus predictable MUA for deep vs. superficial layers. Mean +/- SEM across MUAs in superficial (N= 575, 215, 397 for areas V4, 7A, and PFC) vs. deep layers (N= 615, 248, 468 for areas V4, 7A, and PFC). Red asterisk denotes significant differences.

Neurophysiological analysis focused on the one second of fixation before the sample (pre-sample window), the one second of sample presentation (sample window). During the pre-sample window, monkeys could have an expectation of the forthcoming sample during predictable blocks. Once the sample appeared, predictions about the sample were confirmed or violated.

### Neuronal spiking was greater to unpredictable than predictable stimuli

During the sample window, spike rates were higher if the sample was unpredictable compared to when it was predictable. The black lines in Figure 2A-E show the average multi-unit activity (MUA) for unpredictable minus predictable blocks (see Methods for data pre-processing steps; for MUA response to unpredictable vs. predictable without subtraction, see Supplemental Fig. 1). Positive numbers mean more spiking during unpredictable than predictable blocks. MUA in all areas showed greater spiking (Figure 2A-E, black bars, cluster-based randomization testing, p<0.05) to unpredictable than predictable samples. Further, analysis of thresholded single units (see Methods) confirmed that spikes also carried more information about the sample when it was unpredicted in the sample window (p<0.05 cluster-based randomization test, Figure 2F). V4 spiking carried more sample information than other areas (Supplemental Figure 3A, Wilcoxon rank sum test, V4 vs. all individual areas, all comparisons p<0.01). Further, the increase in spiking to unpredictable samples was stronger in superficial than deep layers (Figure 2G) in area V4 but not in 7A or PFC (Wilcoxon rank sum test for MUA difference, unpredicted minus predicted, in superficial vs. deep at 0.1-0.5s post cue onset, p<0.05).

As expected given randomly drawn samples during unpredictable blocks, spiking during the pre-sample window carried no information about the forthcoming sample. They did during predictable blocks (p<0.05 cluster-based randomization test, Figure 2F). All areas carried significant pre-sample information. PFC carried more information about the forthcoming sample than area V4 (Supplemental Figure 3B, Wilcoxon rank sum test, V4 vs. PFC, p< 0.0001).

### Predictability changed the theta/beta/gamma LFP power balance

Sample predictability had different effects on different oscillatory bands/layers/areas. For each frequency, we calculated percent change in LFP power for unpredictable vs. predictable blocks (Figure 3A-E, black lines). During the sample window, gamma-band power (~40-90 Hz) was higher to unpredictable samples in all areas. In V4 and FEF, theta-band power (~2-6Hz) was higher to unpredictable than predictable sample (black bars in Figure 3A-E, P<0.05, cluster based randomization test). The alpha/beta band (8-30 Hz) showed the opposite effect (Figure 3). It was generally higher to predictable than unpredictable samples, the one exception was posterior parietal area 7A where there was higher power in a high-beta band (~20-27 Hz) during unpredictable samples (Figure 3C – but note that in 7A power in a lower frequency band, 6-14 Hz, was higher for predictable cuing). All areas with laminar recordings (V4, 7A, PFC) showed a greater increase in superficial-layer than deep-layer gamma power during unpredictable samples (Figure 3F, Wilcoxon rank sum test comparing power modulation in superficial vs. deep layers, P<0.05) as did theta in area V4 (Figure 3H). The PFC showed a greater increase in superficial-layer than deep alpha/beta during predictable samples (Figure 3G, Wilcoxon rank sum test, P<1E-4). Area 7A showed a greater increase in superficial-layer than deep beta during unpredictable samples (Figure 3G, Wilcoxon rank sum test, P<1E-4). In, general, the sites with strong MUA modulation (unpredictable > predictable) were also sites with strong LFP gamma power modulation (Supplemental Figure 2). The relationship between MUA and LFP gamma power modulation was consistently strongest in superficial as compared to deep layers (Supplemental Figure 2B, Wilcoxon rank sum test comparing correlation between MUA and gamma power modulation in superficial vs. deep layers, P<1E-4 for all areas).

**Figure 3.**
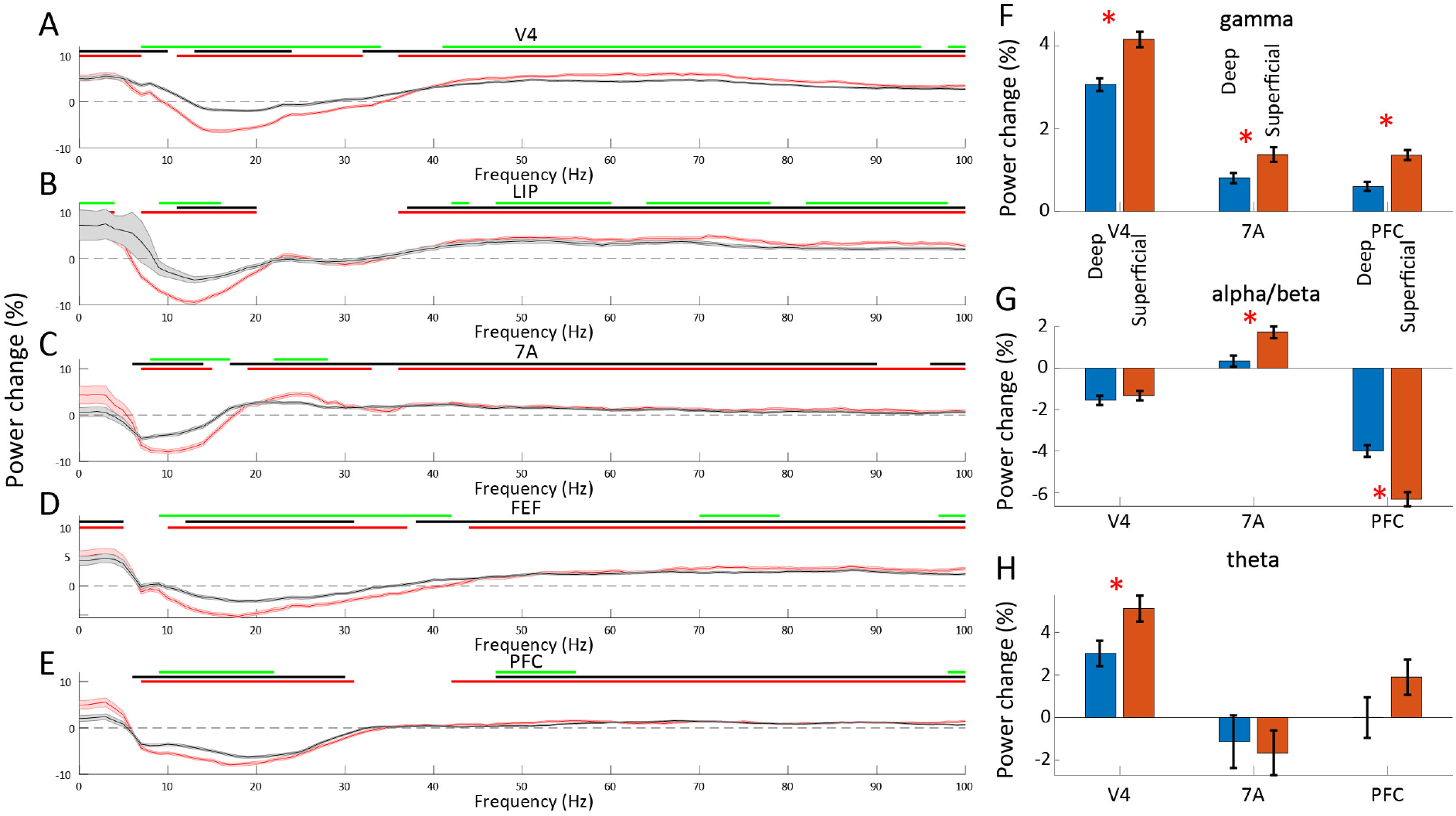
LFP power for unpredictable vs. predictable samples. A-E, Modulation of Local Field Potential (LFP) power from 0-100 Hz during the 1 second sample processing period. Black lines: unpredictable (trials 11-50 in an unpredictable block) minus predictable (trials 11-50 of a predictable block) LFP relative to sample onset (time 0). Red lines: violation (trials 1-10 in an unpredictable block when it is preceded by a predictable block) minus predictable (trials 11-50 of a predictable block) LFP. Mean across all available LFPs per area (N= 1,483, 666, 650, 793, 1,358 for V4, LIP, 7A, FEF, PFC), +/- 1SEM across LFPs. Horizontal bars denote significance at p< 0.05 for unpredictable vs. predictable (black), violation vs. predictable (red), and violation vs. unpredictable (green). F-H, Percent change in LFP power for unpredictable vs. predictable samples for deep vs. superficial layers in different bands. F, gamma-band (40-90 Hz), G, alpha/beta-band (8-30 Hz), H, theta-band (2-6 Hz). Mean +/- SEM across LFPs in superficial (N= 564, 227, 410 for areas V4, 7A, and PFC) vs. deep layers (N= 586, 245, 480 for areas V4, 7A, and PFC). Red asterisk denotes significant (p<0.05) differences.

The increase in gamma and theta tended to be greater in V4 compared to higher areas (Supplemental Figure 3C and E). V4 showed the greatest increase in gamma and theta during unpredictable samples and PFC/7A the smallest. In contrast, higher-order cortical areas PFC and LIP had stronger alpha/beta power to predictable samples than V4 (all comparisons, Wilcoxon rank sum test, p<1E-4), the opposite of gamma.

Sample predictability also modulated LFP power during the pre-sample window but the effects were weaker and sparser. There was greater gamma/high-beta power during unpredictable than predictable blocks in V4, LIP, and 7A, and reduced alpha/beta power in V4, 7A and FEF (p<0.05, Supplemental Figure 4A-E). During the pre-sample window, power modulation did not differ between superficial vs. deep layers (all comparisons, P>0.05).

### Violation of predictions

During predictable blocks, a strong expectation of a specific sample object could build. Then, when there was a switch to an unpredictable block that expectation was violated for at least the first few trials, potentially resulting in prediction errors. We examined the LFP power and MUA during the first 10 “violation” trials of unpredicted blocks. During the sample window, there were increases in MUA and gamma power to violation trials vs predictable samples (Figure 2A-E and 3A-E, red lines) in all areas (Figure 2A-E and 3A-E, red lines show significant differences between violation vs. predictable trials, p<0.05, cluster-based randomization test). The greater increase in alpha/beta power to predictable then violation trials was also significant in all areas (Figure 3A-E, red bars, p<0.05, cluster-based randomization test).

Violation trials showed also stronger effects than the remainder of the “non-violation” trials of the unpredictable block (trials 11-50). In almost all areas, spiking activity and gamma power was higher to violation than unpredictable samples and alpha/beta power was more suppressed (green bars in Figure 2A-E and 3A-E, p<0.05, cluster-based randomization statistics). The exception was gamma in 7A (green bars in Figure 3C, p<0.05, cluster-based randomization). Area 7A alone showed a 22-28 Hz beta enhancement on the violation trials relative to non-violation trials of the unpredictable block. These analyses indicate that violations are a special case of unpredictability and lead to even more enhanced neural spiking and LFP power modulation. Therefore, for all further analyses, violations and unpredictable trials were combined (unpredictable trials) and compared to predictable trials.

### Pathway specificity of LFP power modulation

Predictive Routing posits that effects of predictability should be stimulus/pathway-specific. In other words, the modulation of power should be for strongest for the neural circuits that process the specific stimulus that is being predicted. We addressed this in V4 because it showed the strongest selectivity for the sample objects (Supplemental Figure 3A).

We first analyzed each V4 site’s MUA activity for sample specificity. For each V4 site, the sample that drove the strongest MUA activity was defined as the “preferred” sample, and the sample with the least MUA activity as the “non-preferred” sample. For both the sample and pre-sample window, we calculated differences in power during unpredictable vs. predictable blocks for the preferred and nonpreferred sample separately. If preparatory modulation is pathway specific, it should be greater for the preferred sample and weak or non-existent for the non-preferred sample.

This was indeed the case. During the sample window, LFP gamma power and MUA modulation was stronger (unpredicted > predicted) to the preferred stimulus in superficial cortical layers (p<0.01, Wilcoxon rank sum test comparing each site’s preferred vs. non-preferred power modulation) but not in deep (Figure 4A/B for LFP gamma and Figure 4G/H for MUA). Alpha/beta power modulation (predicted > unpredicted) was stronger to the preferred stimulus in deep cortical layers (Figure 4C, p<1E-3, Wilcoxon rank sum test comparing each site’s preferred vs. non-preferred power modulation) and superficial (Figure 4D, p<0.05, Wilcoxon rank sum test). Also, beta power for the non-preferred stimulus was not modulated at all by predictability (Figure 4C/D, mean power modulation +/- SEM overlaps with zero). Theta power modulation (unpredicted > predicted) was also marginally stronger at sites that preferred that sample for both deep and superficial cortical layers (p=0.056, Wilcoxon rank sum test comparing each site’s preferred vs. non-preferred power modulation, Figure 4E and F). Similar patterns held during the pre-sample window, but with weaker modulation. Theta was not selective for the preferred vs. non-preferred samples in the pre-sample window, but both alpha/beta and gamma were (p<0.05, Wilcoxon rank sum test comparing each site’s preferred vs. non-preferred power modulation, Supplemental Figure 4F-K).

**Figure 4.**
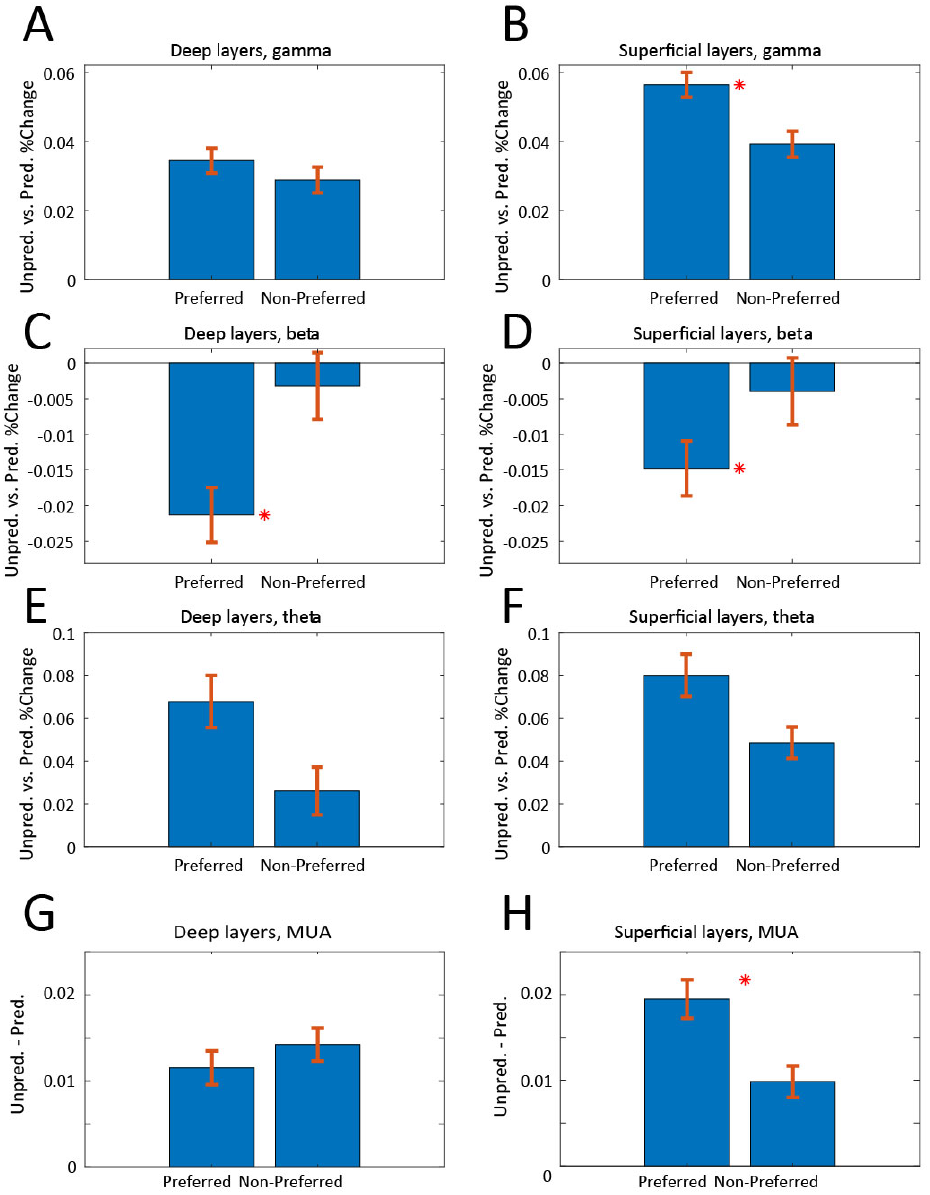
Pathway specificity of LFP power and MUA modulation. A-F, Unpredicted vs. predicted percent change in LFP power for the preferred and non-preferred stimulus. A, deep layers in the gamma-band (40-90 Hz), B, superficial layers in the gamma-band band, C, deep layers in the alpha/beta band (8-30 Hz), D, superficial layers in the alpha/beta band, E, Deep layers in the theta-band (2-6 Hz), F, superficial layers in the theta-band. G, MUA modulation of deep cortical layers in area V4, unpredictable – predictable during the sample period, H, same as G, but for superficial layers. Mean +/- SEM. Red asterisk denotes significant (p<0.05) differences.

### Predictability modulates inter-area coherence

Coherence from 1-100 Hz was calculated between bipolar-derivations of LFPs across all combinations of areas (see Methods). Modulation of coherence by predictability (Unpredictable minus Predictable, in z-score units, N=20,384 inter-areal bipolar site pairs) between one example pair of areas, V4 and FEF, is shown in Figure 5A (all area pairs are shown in Supplemental Figure 5). There was greater coherence (positive values) during unpredictable samples in the gamma and theta bands (see red bars, indicating significance, p<0.05, corrected for multiple comparisons, cluster-based randomization). By contrast, there was greater alpha/beta band coherence to predictable sample (negative numbers, blue bars, Figure 5A, p<0.05, cluster-based randomization).

**Figure 5.**
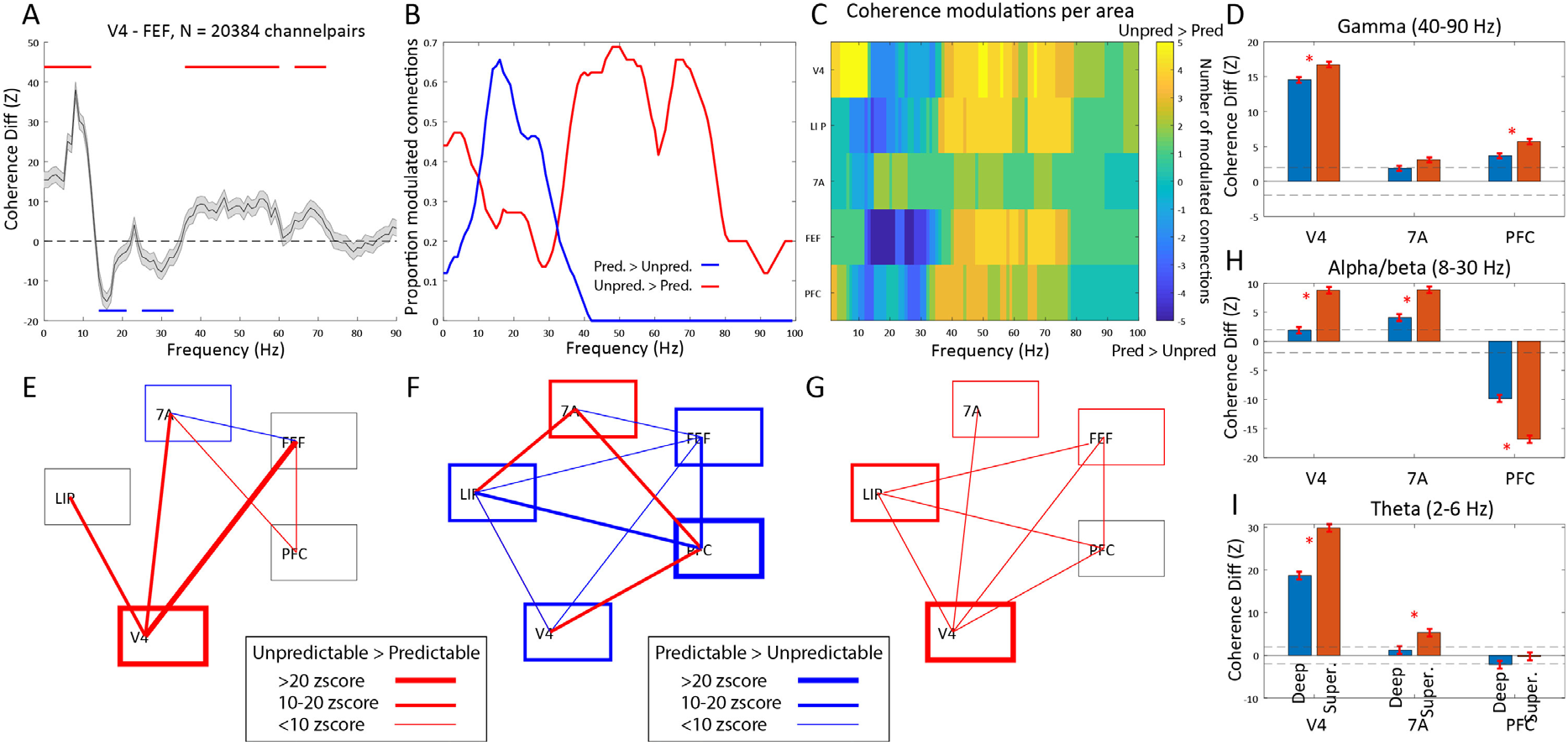
Coherence networks for predicted vs. unpredicted sample processing. A. V4-FEF coherence z-score difference (unpredictable minus predictable) across all inter-area site pairs, N=20,384. Horizontal bars indicate significant (p<0.05, corrected for multiple comparisons) differences for unpredictable > predictable (red bars) and unpredictable < predictable (blue bars).B. Across all possible pairs of cortical areas (N=25, counting within-area connections), the percentage of significantly modulated connections (red: unpredictable > predictable, blue: unpredictable < predictable). C. The number of modulated connections per area as a function of frequency. D, H, I, Coherence z-score difference (unpredictable minus predictable) between each area and the rest, separately for superficial coherence to rest (red bars) and deep coherence to rest (blue bars). Mean +/- SEM, asterisks denote significant differences between layers. E,F,G, Within and inter-area coherence modulations by band, for the theta (2-6 Hz), alpha/beta (8-30 Hz), and gamma (40-90 Hz) bands. The strength of modulation is represented by line thickness (see legend). Edges represent inter-areal coherence modulation. Box outlines represent within-area coherence

Similar effects were seen across the entire network of areas (Figure 5B shows the percent modulated connections across the network, total number of possible connections is 15). Both gamma (40-90 Hz) and theta (2-6Hz) coherence were higher during an unpredictable than predictable samples (Figure 5B, red line) while alpha/beta coherence showed the opposite (Figure 5B, blue line). Figure 5C shows effects of coherence for each area to the rest, with colors representing the number of significantly modulated connections (p<0.01, cluster-based randomization). The network of significantly modulated coherence links is shown in Figure 5E-G. Red indicates greater coherence during unpredictable samples and blue greater coherence during predictable samples with line thickness indicating the strength of effect in z-score units (see Methods). The boxes indicate within-area coherence. Every area showed some modulation of coherence by sample predictability (Figure 5E-G). There was more gamma (and theta, with one exception) coherence during predictable than unpredictable samples and more alpha/beta coherence during predictable than unpredictable samples. But note that as in the power analysis (see above), area 7A was an outlier in the alpha/beta band, showing greater within- and across-area coherence to unpredictable samples (Figure 5F).

There were also differences in coherence between layers. In the gamma band, the superficial layers of V4 and PFC showed a greater increase in coherence during unpredictable than predictable samples (positive numbers, Figure 5D) as did superficial-layer V4 theta coherence (Figure 5I, p<0.05, Wilcoxon rank sum test, comparing all coherence modulations in superficial vs. deep layers). There was also a greater increase in superficial-layer alpha/beta coherence between each area and the network but for V4 and 7A, it was during unpredictable samples while for the PFC it was during predictable samples (Figure 5H p<0.05, Wilcoxon rank sum test, comparing all coherence modulations in superficial vs. deep layers).

During the pre-sample window, coherence was also modulated by predictability. Figure 6A shows the percentage of coherence links modulated by sample predictability with greater coherence during unpredictable blocks shown in red and predictable blocks shown in blue. These pre-sample modulations (Figure 6A/B) were strikingly different during the sample (see above and Figure 5B/C). First, modulation of gamma-band coherence was very low pre-sample, as expected, because of the lack of bottom-up sensory input. Second, there was a flip in the sign of coherence modulations for theta and alpha/beta. In theta, coherence was greater during predictable than unpredictable Blocks during the pre-sample (Figure 6A) while the opposite was seen during the sample (Figure 5B). Likewise, pre-sample alpha/beta band coherence showed a greater increase to the rest of the network during unpredictable relative to predictable blocks (Figure 6A), the opposite to that seen during the sample (Figure 5B).

**Figure 6.**
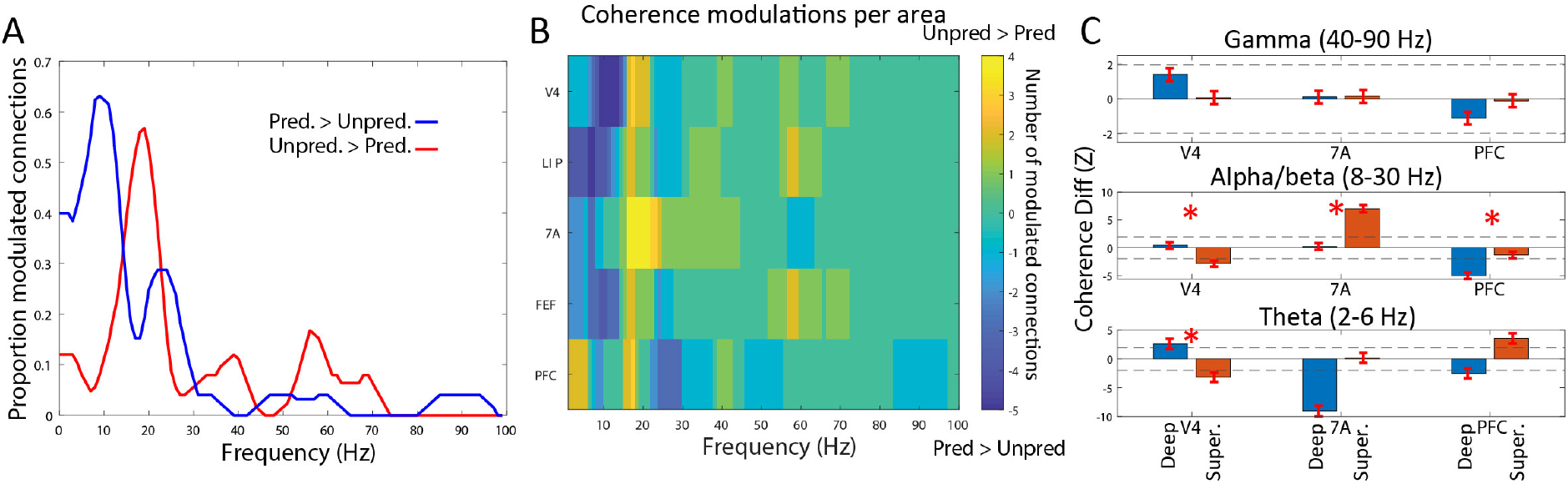
Coherence networks in the pre-sample window. A. Same as Figure 5B, but for the pre-sample window. B. Same as Figure 5C, but for the pre-sample window. C. Same as Figure 6 E,F,G, but for the pre-sample window.

There were also layer differences during the pre-sample period. PFC showed significantly more alpha/beta coherence to the rest of the network in deep than superficial layers during predictable than unpredictable Blocks (Figure 6C middle subpanel, p<0.05, Wilcoxon rank sum test comparing all coherence modulations in superficial vs. deep layers). By contrast, superficial-layer V4 showed a greater increase in network theta coherence during predictable blocks (Figure 6C lower subpanel, p<1E-8, Wilcoxon rank sum test comparing all coherence modulations in superficial vs. deep layers). Finally, area 7A showed a greater increase in superficial than deep alpha/beta coherence during unpredictable blocks (Figure 6C middle subpanel, 1E-9, Wilcoxon rank sum test comparing all coherence modulations in superficial vs. deep layers). No pre-sample gamma coherence differences exceeded +/- 2 Z-score units of difference between predictable vs. unpredictable conditions.

### Directed network interactions

We next examined directed interactions between areas. We used Granger Causality (GC), which separately measures the impact of area A to B vs. B to A at each frequency from 1-100Hz (Ding et al., 2006). To assess feedforward vs feedback flow, we assumed the following cortical hierarchy (from lower to higher): V4, LIP, 7A, FEF, PFC (Felleman and Van Essen, 1991). We first focused on the sample window. Figure 7A shows the proportion of significantly modulated connections (cluster-based randomization test, p<0.01) for both feedforward (solid lines) and feedback (dotted lines) directions (Modulation of Granger causality for all individual area pairs is shown in Supplemental Figure 6). The red line indicates more modulation to unpredictable than predictable samples whereas the blue line shows the opposite (cluster-based randomization test, p<0.01). Figure 7D shows the sums of modulated connections per area, as a function of whether connections into and out of the area were feedforward or feedback.

**Figure 7.**
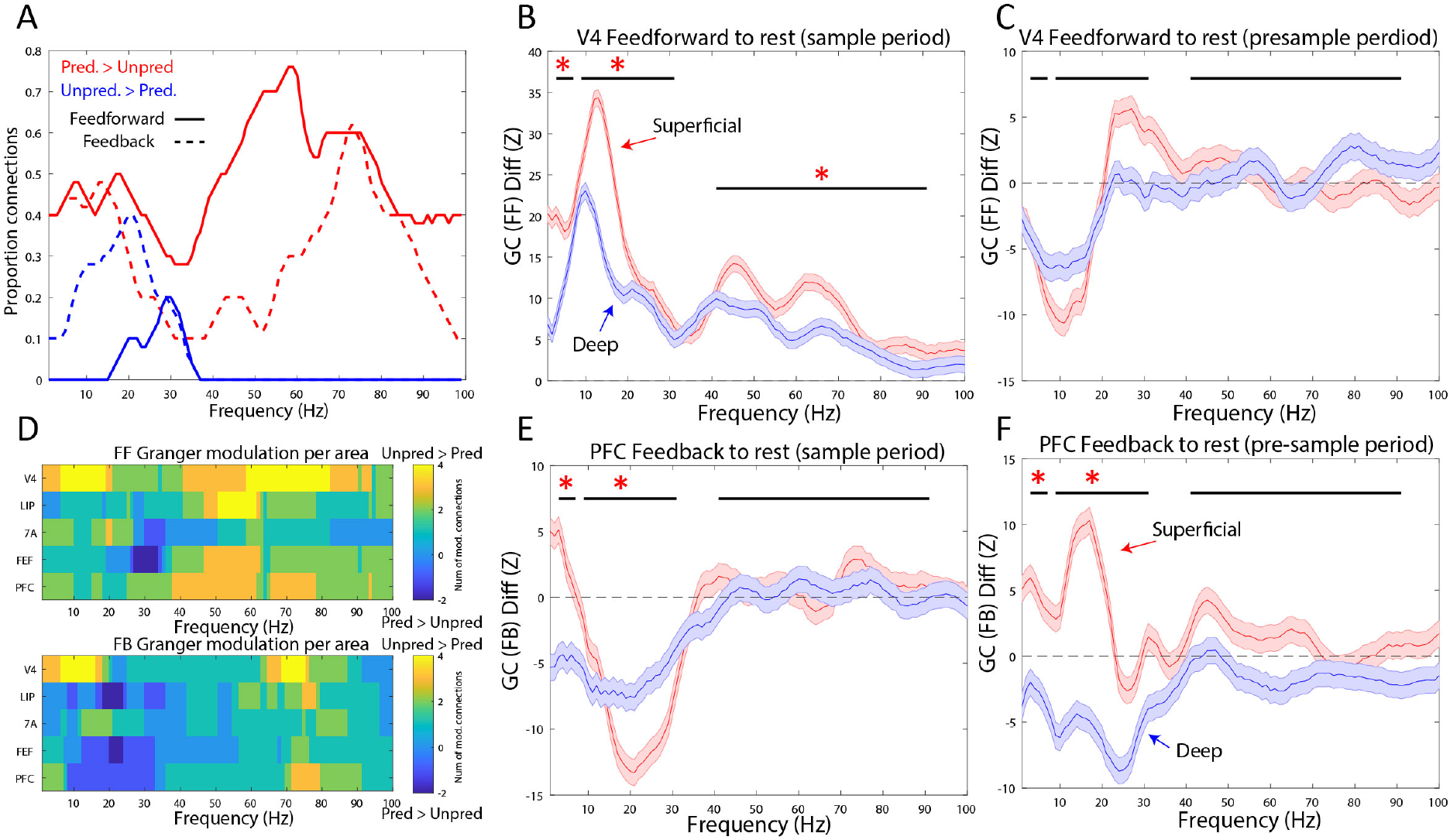
Granger causal networks for predicted vs. unpredicted cue and pre-cue processing. A. Percentage of inter-areal functional connections with significant (p<0.05) task modulation (red lines: unpredictable > predictable, blue lines: unpredictable < predictable), separately for feedforward (solid lines) and feedback (dotted lines). B. Granger causal z-score difference from V4 to other areas during the sample period for superficial (red lines) vs. deep (blue lines). Red asterisks denote frequency ranges with significant laminar differences. Mean +/- SEM. C. Same as B, but for the pre-sample period. D. Number of feedforward functional connections (upper subpanel) and feedback functional connections (lower subpanel) modulated by frequency (yellow colors represent number of functional connections with more unpredictable > predictable, blue lines represent number of functional connections with more unpredictable < predictable). E and F, same as B and C, but for feedback connections from PFC to other areas. Mean +/- SEM.

The strength and sign of modulation of GC by predictability depended on frequency and directionality. In the feedforward direction, unpredictable > predictable GC modulation peaked in the theta and gamma frequency ranges (Figure 7A, solid red lines). Although the feedback direction was also positively modulated (Figure 7A, dotted red lines), the proportion of modulated connections was lower than feedforward gamma (Chi-squared test for proportion of positive modulation across the gamma band, 40-90 Hz, in the feedforward vs. feedback directions, 0.57 vs. 0.33, p< 1E-13). In the theta band (2-6 Hz), an approximately equal proportion of both feedforward and feedback connections had positive task modulation (unpredicted > predicted GC).

Notably, virtually all directed functional connections with greater GC during predicted than unpredicted GC) occurred in the alpha/beta band (blue lines in Figure 7A). They were mostly feedback connections. The proportion of feedback direction GC that was greater for predictable than unpredictable samples was higher than those in feedforward direction for both alpha/beta (Chi-squared test for proportion of negative modulation across the alpha/beta band, 8-30 Hz, in the feedforward vs. feedback directions, 0.07 vs. 0.30, p< 1E-9) and theta (Chi-squared test for proportion of negative modulation across the theta-band in the feedforward vs. feedback directions, 0.00 vs. 0.1, p< 0.05). There was no predictable > unpredictable GC modulation in the gamma-band in either direction. In short, feedforward functional connections are enhanced during unpredictable samples, especially in the gamma range whereas feedback functional connections were enhanced during predictable samples in the alpha/beta and theta frequencies.

To determine the layer specificity of these effects, we focused on the two areas at the bottom and top of the hierarchy: V4 and PFC. The rationale was that GC interactions from V4 to the other areas are all feedforward, and interactions from PFC to the other areas are all feedback. The modulation of these feedforward and feedback interactions by layer is shown in Figure 7B (for V4) and 7E (for PFC) during the sample window, and for the same areas during the pre-sample window in Figure 7C and F.

During the sample window, feedforward connections from V4 to the rest of the areas were greater for unpredictable samples and this modulation was greater in superficial layers compared to deep in theta, alpha/beta and gamma frequency bands (Figure 7B, Wilcoxon rank sum test comparing modulations for all feedforward electrode channel pairs in superficial vs. deep layers, p<1E-4 for theta, p<1E-8 for alpha/beta, and p<1E-4 for gamma). By contrast, in PFC during the sample, feedback GC was greater for predictable samples, especially in the alpha/beta band (Figure 7E). This effect was stronger in superficial than deep layers (Figure 7E, Wilcoxon rank sum test comparing task modulations for all feedback channel pairs in superficial vs. deep, p<1E-12 for the alpha/beta band). In the theta-band, PFC feedback GC in deep layers was significantly stronger for predictable samples compared to superficial layers (Figure 7E, Wilcoxon rank sum test comparing task modulations for all feedback channel pairs in superficial vs. deep, p<1E-5 for the theta band).

During the pre-sample window, PFC feedback interactions were different depending on whether they arose from superficial vs. deep layers. Superficial-layer theta and alpha/beta PFC feedback was greater during unpredicted blocks but deep-layer PFC feedback was greater during predicted blocks. The differences between layers was significant for both theta and alpha/beta (Figure 7F, Wilcoxon rank sum test comparing modulations for all feedback channel pairs in superficial vs. deep layers, p<0.001 for the theta band, p<1E-10 for the alpha/beta band). In contrast, there were no laminar differences in the modulations of feedforward interactions between V4 and the other areas by Unpredictable vs Predictable cue blocks (Figure 7C).

## Discussion

### Relation to predictive coding models

Many predictive coding models share common elements (Figure 8A-C). Prediction (PD) units anticipate forthcoming sensory inputs. They inhibit prediction error (PE) units when inputs match predictions. A mismatch due to an unpredictable input disinhibits the PE units. They feedforward the unpredicted input which update the internal models that generate the predictions.

**Figure 8.**
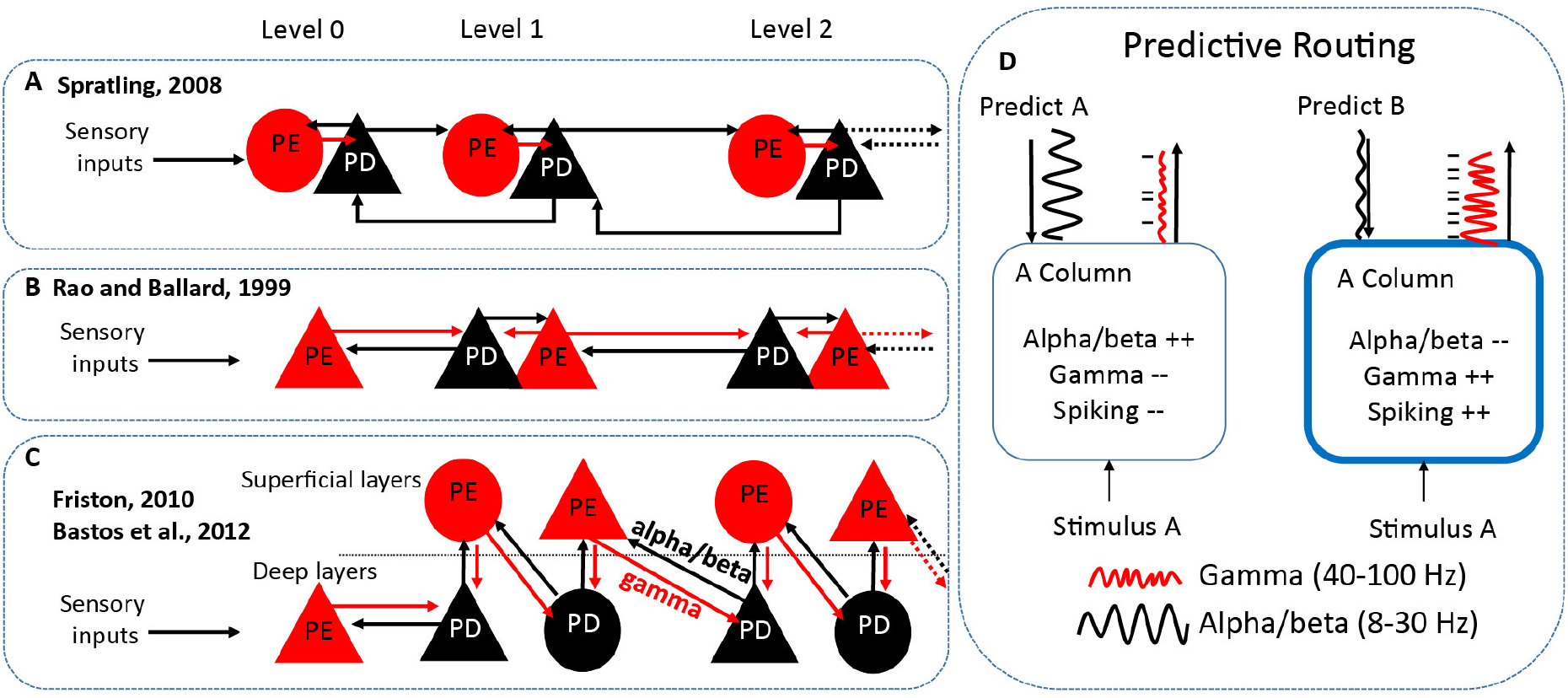
Models of predictive coding and routing. Architecture of current predictive coding models and predictive routing. A-C, current predictive coding implementations. Red circles/triangles denote prediction error (PE) units, black circles/triangles denote prediction (PD) units, and arrows their targets. Intrinsically projecting cells (within-area) are round, extrinsically projecting units (inter-areal projections that traverse hierarchical levels) are triangles. Further levels are implied, denoted by dashed arrows. Model A (Spratling et al., 2008) hypothesizes that PE signals stay local within an area, and the PD is signaled between areas in feedforward and feedback directions. Model B (Rao and Ballard, 1999) hypothesizes that PE is sent feedforward and PD is sent feedback. Model C (Friston et al., 2010, Bastos et al., 2012) hypothesizes that cells in superficial layers send feedforward PEs (at gamma) and cells in deep layers send feedback PDs (at alpha/beta). Models A-C all hypothesize al specialized circuit for error processing, wherein PD units inhibit PE units. D, Predictive Routing model. (left subpanel) Sensory cortex is dynamically prepared to process its preferred stimulus, stimulus A, by feedback alpha/beta. Enhanced alpha/beta functionally inhibits processing of stimulus A by reducing spiking and gamma, reducing feedforward outputs. (right subpanel) In the absence of a prediction for stimulus A, there is less feedback alpha/beta to the A column. The A column is more excitable, and responds to stimulus A with more gamma and spiking and enhanced feedforward output (a “prediction error”).

Models differ on the details of the implementation in the brain. Some models (Spratling, 2008, Figure 8A) propose that prediction error signals act locally in each cortical area to update models. Predictions flow between areas in both feedforward and feedback directions. Other models (Rao and Ballard, 1999, Friston, 2010, Bastos et al., 2012) instead suggest that predictions come from higher cortical areas that act on lower cortical (sensory) areas to gate the feeding forward of prediction errors (Figure 8C/D). “Rhythm-based” models (Figure 8C) suggest that superficial cortical layers (layers 2 and 3) feedforward prediction errors using gamma. Deep-layer cortex (layers 5 and 6) feedback predictions using alpha/beta (Arnal and Giraud, 2012; Bastos et al., 2012).

Our results are more consistent with the rhythm-based models. We showed differences between areas with higher areas contributing more to prediction, and rhythmic and laminar based differences between areas. They also suggest an update: Predictive Routing.

### Predictive Routing

Alpha/beta and gamma have properties that suggest a general role in gating and control. They are common and anticorrelated across cortex. Gamma power is high during sensory inputs; alpha/beta power is high when they are ignored. In visual cortex, gamma power is high and alpha low during sensory stimulation. When a stimulus needs to be filtered or ignored, gamma power is low and alpha is high (Bauer et al., 2006; Buffalo et al., 2011; Fries et al., 2001; Jokisch and Jensen, 2007). Further, alpha/beta oscillations in deep cortical layers are anticorrelated with superficial-layer gamma associated with spiking carrying sensory inputs (Bastos et al., 2018; Lundqvist et al., 2016). The balance between alpha/beta and gamma reflect the encoding, maintenance, and read-out of working memory (Lundqvist et al., 2018). We have lumped together alpha and beta together in this paper because they are both functionally inhibitory (negatively correlated with spiking, see Supplemental Figure 2). The general idea is that top-down signals are fed back through alpha/beta in deep cortical layers. They inhibit and thus gate the expression of gamma in superficial layers that help feed forward and maintain the spiking carrying sensory inputs.

Extrapolating from the rhythm-based models of predictive coding (Arnal and Giraud, 2012; Bastos et al., 2012), this suggests Predictive Routing (PR, Figure 8D). In PR, there are not specialized circuits that compute prediction errors and send them feedforward. Rather, PR uses the same cortical circuitry used for other functions (sensory processing, attention, maintenance/control of working memory, etc.). Predictions act by alpha/beta preparation that actively inhibit the specific pathways in sensory cortex that would process the predicted input. As a result, there is less gamma and spiking to predicted inputs (and less feedforward output as a result). In other words, prediction errors do not result from a comparison between predictions and inputs via a specialized circuit. They result from the feed forward passing of unexpected inputs because their pathways have not been prepared (functionally inhibited).

This does not mean that alpha/beta (or gamma) have the exact same roles all over cortex. For example, in prefrontal cortex, beta has been modeled as an inhibitory-excitatory network but with slower time constants than what would produce a gamma (Sherfey et al., 2018). By contrast, modeling suggests that in parietal cortex, there is a distinction between beta-1 (14-20 Hz) and beta-2 (24-30 Hz) (Roopun et al., 2008) and that parietal beta-1 may act as a memory buffer activated by strong cortical inputs that feeds forward violations to the PFC (Gelastopoulos et al., 2019). Indeed, we found in this study, that in parietal cortex (area 7A) beta was unique, beta was functionally excitatory (positively correlated with violations) unlike the other areas. Below, we elaborate on how our results support different models of predictive coding.

Theta-band coherence and Granger causal interactions were stronger during unpredictable stimuli in the sample window. These interactions were strongest between superficial layers of V4 and the other members of the network. Theta-band interactions were not previously proposed as candidates for processing unpredictable stimuli. However, theta is well known to be a slow rhythm in which faster rhythms such as gamma, can nest (Canolty et al., 2006; Herman et al., 2013; Lakatos et al., 2008) and aid long-range communication (Tort et al., 2007). In addition, a previous study identified theta as a carrier for feedforward interactions in the visual system (Bastos et al., 2015). In predictive routing, whatever mechanisms are already in place for feedforward processing are enhanced during unpredictable processing. Therefore, it makes sense that theta (and gamma) from V4 to higher-order cortex is enhanced during unpredicted stimuli, because this reflects an upregulation of the feedforward channel.

### Unpredicted stimuli enhance spiking in superficial layers

At each level of the cortical hierarchy tested, neurons spiked more to, and carried more information about, unpredictable compared to predicted stimuli and even more to violations (an unexpected stimulus when there was a strong prediction). Neurons in superficial cortical layers (L2/3) of V4 showed stronger effects than deep layers. In addition, only superficial layers had unpredicted spiking selective to the predicted stimulus. Superficial layers 2/3 contain the majority of feedforward projecting cells. This laminar specificity suggests that unpredicted inputs are preferentially processed in superficial layers and fed-forward.

This effect could be explained, at least in part, by stimulus-specific adaption (SSA) (Miller et al., 1991). According to SSA, synapses “habituate” when repeatedly activated. Thus, there is greater spiking to non-repeated inputs. However, this does not explain the laminar specificity (in superficial layers, as hypothesized in Bastos et al., 2012, Figure 8C), the enhanced selectivity of superficial layers, or the observed changed in oscillatory dynamics seen at the LFP level.

### Spiking reflects a predictable stimulus

We found that during trial blocks in which the monkeys could predict the upcoming stimulus, all cortical areas we recorded carried information about it before it appeared. PFC contained the most pre-stimulus information. This is more consistent with hierarchical models that propose that generating predictions is primarily a function of higher than sensory cortex.

### Modulation of rhythms by prediction are layer-dependent

In all areas studied, unpredictable stimuli evoked more gamma (40-90 Hz) power and less alpha/beta (8-30 Hz) power compared to predictable stimuli (Area 7A was an exception). These effects were strongest when there was a strong prediction, as opposed to when stimuli were highly unpredictable. This effect and the positive correlation between gamma and spiking (Supplemental Figure 2) were stronger in superficial cortical layers than deep cortical layers in all areas tested. There was generally more alpha/beta power to predictable than unpredictable stimuli but layer differences varied by area This generally supports rhythm-based models positing that gamma helps transmit prediction errors and alpha/beta helps transmit predictions.

The increases in gamma power (and spiking) with unpredicted stimuli and the increased alpha/beta with predicted stimuli were stimulus-specific. Effects were larger at recoding sites where spiking “preferred” (was greater to) the specific stimulus that was predicted. For LFP gamma and spiking, this selectivity occurred only in superficial layers. The specificity is a central feature of Predictive Routing, where alpha/beta inhibits the pathways that process the specific predicted stimulus and the increased gamma/spiking occurs because the pathways processing other stimuli were not inhibited.

### Networks and directionality

Both coherence and Granger Causality analysis showed that rhythmic interactions were modulated by stimulus predictability at several frequencies. Gamma-band coherence within and between areas was higher with unpredictable than predictable stimuli. This effect was largest for coherence between superficial layers of areas V4 and PFC. Granger Causality analysis further showed that the increase in gamma band coherence with unpredictable stimuli was stronger in the feedforward than feedback direction. In V4, this was more prominent in superficial layers. There was overall greater alpha/beta coherence with predictable stimuli. The strongest effects involved PFC and were stronger in the feedback compared to feedforward direction. In the pre-sample period, the enhanced Granger Causality during predictable stimuli was strongest between deep layers of PFC to the rest of the network. These results are in line with hierarchical and rhythms models where (gamma-based) prediction errors primarily feed forward flow up the cortical hierarchy and (alpha/beta-based) predictions flow down the cortical hierarchy. They suggest that modulation of inter-areal synchronization at distinct frequencies is a central mechanism in communicating specific (predicted vs. unpredicted) information (Fries, 2015; Womelsdorf et al., 2007, 2014a, 2014b). In addition, prefrontal control over behavior is thought to be mediated by dynamic patterns of neuronal functional connectivity (Crowe et al., 2013). In future work, we will mechanistically explore how the inter-areal theta, alpha/beta, and gamma synchronization task-related changes we have observed compare to the neuronal code expressed by each area.

### Summary

Our results suggest a hierarchical, layer, and frequency-specific model for predictive coding that we term “Predictive Routing” (Figure 8D). Unpredictable stimuli evoked stronger feedforward-superficial-layer gamma (and theta), especially when they violated a previous prediction; the hallmark of a prediction error signal. Superficial-layer parietal area 7A high beta also signaled violations in both feedforward and feedback directions, which could engage working memory update mechanisms to process and hold unpredicted information online. Coherence and feedback connectivity were enhanced in the alpha/beta band when a stimulus was predictable. In the pre-sample period this enhanced feedback connectivity during predictable stimuli originated in deep layers of PFC. The modulatory effects of stimulus predictability on alpha/beta and on gamma/spiking modulation was strongest at the sensory cortical sites that preferred the predicted stimulus. Gamma and spiking in sensory cortex were only selective to the predictive stimulus in superficial layers. Together, these results suggest that predictive coding may stem from rhythmic interactions between lower frequency rhythms in deep cortical layers that signal predictions and inhibit the superficial-layer gamma and spiking in the sensory pathways that match those predictions.

## Author Contributions

AMB and EKM designed the study and wrote the manuscript with input from the other authors. AMB trained animals, designed implants/recording approach, performed surgeries, collected and analyzed data. EKM supervised the study and acquired funding. NK co-supervised the analysis and formulated the predictive routing model, with input from AMB and EK. ML assisted with recordings and trainings. ASW assisted with data post-processing and spike sorting.

## Acknowledgements

We would like to thank Scott Brincat for assistance with surgeries and data pre-processing, and Morteza Moazami and Jefferson Roy for assistance with surgeries and animal training. We also thank the MIT veterinary staff and animal caretakers for their excellent support. We also thank Jaan Aru and Bruno Gomes for comments to the manuscript. This work was supported by National Institutes of Mental Health Grant R37MH087027 and 5K99MH116100-02, Office of Naval Research Multidisciplinary University Research Initiatives Grant N00014-16-1-2832, and the MIT Picower Institute Faculty Innovation Fund.

## STAR Methods

### Lead Contact and Materials Availability

The Lead Contact for this study is Earl Miller. Requests for materials should be directed to the Lead Contact. This study did not generate novel reagents.

### Experimental Model and Subject Details

Two adult rhesus macaques (*macaca mulatta*) were used in this study (Monkey S: 6 years old, 5.0 kg and monkey L: 17 years old, 10.5 kg). Both animals were pair-housed on 12-hr day/night cycles and maintained in a temperature-controlled environment (80°F). All procedures were approved by the MIT IACUC and followed the guidelines of the MIT Animal Care and Use Committee and the US National Institutes of Health.

### Methods Details

#### Behavioral Training and Task

Monkeys were trained to sit comfortably in a primate chair inside a sound attenuating behavioral testing booth. They were seated 50 cm away from a LCD monitor with 144Hz refresh rate (ASUS, Taiwan). Using positive reinforcement, we trained monkeys to perform a visual search task (Figure 1A). Monkeys fixated a point at the center of the screen (fixation window radius: 2-3 visual degrees) for a duration of 1s, were presented with one of three cue objects for a duration of 1s, and were required to maintain fixation over a delay (between 0.5-1.2s). A search array then appeared that consisted of the cued item together with either one or two distractors presented at the same eccentricity (3-8 degrees), but different visual quadrants as the cued object. The position of the cued object and the distractors were always randomly chosen. Monkeys were rewarded with a few drops of diluted juice if they performed a saccade toward the cued item. Behavioral performance was high for each of the monkeys (monkey S: 77% over 41 sessions, monkey L: 75% over 30 sessions). Monkeys were trained on this task using a library of 22 sample images. For recordings, we used a subset of these images (12), choosing a total of 3 per session. Most sessions (65 out of 71) used the three objects depicted in Figure 1: an orange, a green block, and a blue car.

To manipulate prediction, the task was performed either with unpredictable or predictable cuing. During unpredictable cuing, samples were randomly drawn on each trial. In block cuing/sampling, the sample was held constant for the duration of the block. The trial-by-trial and blocked modes each lasted for 50 trials before switching block modes. The starting order was randomized over sessions. The task design is schematized in Figure 1A.

#### Surgical Procedures

All procedures were performed in a sterile surgical suite, with animals under full general anesthesia. Animals were first anesthetized with ketamine and then intubated. They were maintained in a stable plane of anesthesia with sevofluorane. After each procedure, animals received analgesic and antibiotic medications. Three surgical procedures were performed per monkey. First, a titanium head post was fixed to the posterior part of the cranium with titanium screws. The head post was allowed to integrate into the bone for at least 8 months prior to the next surgery. Second, a custom-machined Carbon PEEK chamber system with three recording wells (placed over prefrontal, parietal, and visual cortex) was affixed to the cranium, also with titanium screws. After a one-month period, a third procedure was performed, in which three craniotomies ranging in circular diameter between 10-16 mm^2 were opened inside each recording well.

#### Neurophysiological Recordings

All of the data were recorded through Blackrock headstages (Blackrock Cereplex M, Salt Lake City, UT), sampled at 30 kHz, band-passed between 0.3 Hz and 7.5 kHz (1st order Butterworth high-pass and 3rd order Butterworth low-pass), and digitized at a 16-bit, 250 nV/bit. All LFPs were recorded with a low-pass 250 Hz Butterworth filter, sampled at 1 kHz, and AC-coupled.

We implanted the monkeys with a custom-machined Carbon PEEK chamber system with three recording wells placed over visual/temporal, parietal, and frontal cortex. The process for making the chambers was based on design principles outlined previously (Mulliken et al., 2015). Briefly, we first took an anatomical MRI scan (0.5mm^3 voxel size) and/or CT scan to extract the bone and co-register the skull model with the brain tissue. We designed the center of each chamber to overlie the primary recording area of interest and to have an optimal angle for perpendicular recordings relative to the cortical folding. Post-operatively, after the recording chambers were implanted, MRIs were taken with the recording grid in place, filled with water, which created a marker to co-register each possible electrode trajectory with the animal’s anatomy, and to confirm trajectories that were as close to perpendicular as possible.

The areas where we could achieve perpendicular recordings (for laminar sampling) on the overlying gyrus were V4 (Foveal and Parafoveal representations), parietal cortex (area 7A), and prefrontal cortex (area 8A, Ventro and Dorsal lateral prefrontal cortex – VLPFC/DLPFC). The areas where we recorded without laminar alignment (due to their location in sulci) were areas FEF and LIP.

We recorded a total of 71 sessions with laminar probes. In each session, we inserted between 1-3 laminar probes (“U probes” and “V probes” from Plexon, Dallas, TX) into each recording chamber with either 100 or 200 um inter-site spacing and either 16 or 32 total electrodes per probe. This gave a total linear sampling of 3.0-3.1mm on each probe. Between 3-7 probes in total per session were used, with a total channel count ranging between 48-128 electrodes per session. The recording reference was the reinforcement tube, which made metallic contact with the entire length of the probe (total probe length from connector to tip was 70mm). Some U/V probes had noisy channels (average power greater than 2 standard deviations above the mean of all channels, this occurred on less than 5% of all channels), which were interpolated based on nearest neighbors prior to analysis.

#### Lowering procedure and laminar placement of electrodes

Traditionally, studies with laminar probes have used Current Source Density (CSD) mapping to identify the position of layer 4. However, for parietal area 7A, to our knowledge there is no published study using this technique. Therefore, we chose to align our data to the pial surface of the cortex, a technique that has been previously used to separate recording channels into superficial vs. deep layers in monkeys for recordings in parietal and prefrontal cortex (Johnston et al., 2019). This was the most robust laminar alignment metric that could be applied to visual, parietal, and prefrontal cortex with minimal assumptions. The average cortical thickness for the recorded regions was 2.4mm, measured using MRI, and visually measuring the gray matter thickness using Osirix software (Geneva, Switzerland). Superficial layer channels were classified from the top of cortex to a depth of 1.2 mm, and deep layer channels from 1.2 to 2.4mm. This maps approximately onto layers 1-4 for superficial, and 5-6 for deep.

We first punctured the dura using a guide tube. Then we lowered the laminar probes through the guide tube using custom-built drives that advanced with a turn screw system. In order to place the contacts of the laminar probe uniformly through the cortex, spanning from cerebrospinal fluid through the gray matter to the white matter, we used a number of physiologic indicators to guide our electrode placement, as previously described (Bastos et al., 2018). First, the presence of a slow 1-2 Hz signal, a heartbeat artifact, was often found as we pierced the pia mater and just as we entered the gray matter. Second, as the first contacts of the electrode entered the gray matter, the magnitude of the local field potential increased, and single units and/or neural hash became apparent, both audibly and visually with spikes appearing in the online spike threshold crossing. Once the tip of the electrode transitioned into the gray matter, electrodes were lowered slowly an additional ~2.5mm. At this point, we retracted the probe by 200-400 um, and allowed the probe to settle for between one to two hours before beginning the task. We left between 1-3 contacts out of gray matter in the overlying Cerebral Spinal Fluid (CSF).

#### Multi-Unit Activity Extraction and Spike-Sorting

For the analysis of the analog multi-unit activity (MUA) we band-pass filtered the raw, unfiltered, 30kHz sampled data into a wide band between 500-5,000Hz, the power range dominated by spikes. The signal was then low-pass filtered at 250Hz and re-sampled to 1,000 kHz. The advantage of this signal is that it captures all nearby units, including those with low signal to noise ratio that would not be captured with a strict threshold.

For the analysis of thresholded spikes, we first placed an online threshold manually on each recording session to ensure each recording channel captured waveforms with sufficient signal to noise ratio to qualify as neuronal spiking. These were typically placed at between 2-4 standard deviations away from the noise floor. Offline, spike sorting was performed manually using Plexon offline sorter. We projected the waveform shapes into the top 2 or 3 principle components, and sorted each electrode’s threshold crossings into isolatable waveforms. We included these single units into analysis if their average firing rate per trial (between 1.5 seconds pre-sample to 1.5 seconds post-sample) was stable for at least 120 trials. We defined stability with the Matlab function findchangepts().

#### Local Field Potential power, coherence and Granger causality analysis

All analyses were performed with customized MATLAB scripts and with Fieldtrip software (Oostenveld et al., 2011). Bipolar derivation is a recommended pre-step prior to Granger causality and coherence analysis, as the presence of a common reference can lead to spurious results (Trongnetrpunya et al., 2015; Vinck et al., 2015). In addition, bipolar derivation enhances the spatial localization of LFP signals and removes the common reference and any common noise or volume conduction in the signal (Bastos and Schoffelen, 2015). Here, we computed the sample-by-sample bipolar differences by subtracting contacts that at a distance of 400um: next-nearest neighbors for the laminar probe data spaced at 200um between contacts, and next-next-nearest neighbors for the probe data spaced at 100um between contacts.

We then estimated power, coherence, and Granger causality on these bipolar derivations. We estimated power at all frequencies from 0-250 Hz using multitaper spectral estimation (smoothing window of 5Hz, leading to 9 tapers per spectral estimate, using window sizes of 1 second (0 to 1 seconds relative to sample onset is the period of visual stimulation, −1 to 0 seconds relative to sample onset is the pre-stimulus fixation window) per trial. These Fourier coefficients were then used to calculate the Cross-Spectral Density matrix, from which we derived coherence and non-parametric spectral Granger causality (see below).

The computation of Granger causality in the frequency domain requires the estimation of two quantities: the spectral transfer matrix (H(ω)), which is frequency dependent, and the covariance of the model’s residuals (Σ). The spectral transfer matrix defines how power in one channel is transferred to other channels, at each temporal lag. The model’s residuals is not a function of frequency, and defines the amount of variance that is left unexplained by the linear model, H(ω). Traditionally, H(ω) is computed in a parametric (model-based) fashion by first fitting an autoregressive model to the data, and then Fourier transforming the model (Ding et al., 2006). However, it is also possible to compute, H(ω), and thus Granger causality, directly from the spectral transform of the data. In brief, the following fundamental identity holds: H(ω)ΣH(ω)* = S(ω), with S(ω) being the cross-spectral density matrix at frequency ω. Starting from the cross-spectral density matrix (S(ω)) it is possible to factorize the cross-spectral density matrix into a noise covariance matrix (Σ) and spectral transfer matrix (H(ω)) by applying spectral matrix factorization (Dhamala et al., 2008)—which provides the necessary ingredients for calculating Granger causality. The nonparametric estimation of GC has certain advantages over parametric approaches in that it does not require the specification of a particular autoregressive model order.

#### Statistical Testing

We computed whether the MUA, power, coherence, and Granger causality was systematically different between conditions (Predictable vs. Unpredictable, and for a subset of analyses, Predictable vs. Unpredictable vs. Violation trials). To do this, we calculated either the mean difference or percent change for each channel or inter-areal channel-pair of Predictable vs. Unpredictable sampling. We then quantified whether this raw difference or percent change was significant by performing a permutation test (Maris and Oostenveld, 2007). Per channel (for MUA and LFP power) or channel-pair (for coherence and Granger) we randomized the experimental label (Predictable vs. Unpredictable sampling). We performed this randomization 1,000 times. For each randomization, we took the number of consecutive time bins (in the case of MUA) or frequency bins (for power, coherence, and GC) that passed a first level criteria. This first level criteria was that two measures (for example, LFP power when comparing unpredictable vs. predictable cuing) had to differ from one another at significance value of p<0.01, uncorrected, based on a t-test statistic. The maximum positive or negative cluster was determined for each randomization. This step controls for multiple comparisons across neighboring (and possible correlated) time or frequency bins. Finally, these clusters from the randomization distribution were used to determine significance values for the empirically observed clusters, using a p-value of 0.05, corrected for multiple comparisons.

For determining differences between superficial vs. deep layers, we first averaged the corresponding metric (power, coherence, Granger causality) across the particular frequency band (theta, alpha/beta, gamma) for all superficial and deep channels/channelpairs. Then we performed a Wilcoxon rank sum test to determine significant differences across the populations with an alpha at p<0.05.

For determining the difference between feedforward vs. feedback Granger causality modulation, we determined the percentage of modulated inter-areal directed functional connections, integrating all frequencies in the theta, alpha/beta, and gamma frequency bands. We then applied a Chi-squared test for differences in proportion to test whether e.g., feedforward functional connections were more modulated (had a greater proportion) than feedback functional connections.

#### Data and Code Availability

The data and code will be made available by reasonable request by contacting the lead author, Earl Miller.

