## Supplemental Materials for "Layer and rhythm specificity for predictive routing"

**Supplemental Figure 1 | Unpredicted vs. predicted Multi-Unit Activity**

For each area, the average MUA response to an unpredicted cue (red) vs. a predicted cue (blue). Mean +/- SEM across MUAs.

**Supplemental Figure 2 | Correlation between MUA and LFP power by area**

A, Unpredicted minus predicted MUA difference Spearman correlation across time (-1.25 to 1.25 seconds from cue onset) to LFP power at each frequency from 0-100 Hz. Mean Spearman correlation across each area, +/- SEM. B-D, Average Spearman correlation between power and MUA in superficial (right orange subplots) vs. deep (left blue subplots) layers for A, theta (2-6 Hz), B, alpha/beta (8-30 Hz), and C, gamma (40-90 Hz). Mean +/- SEM. Red asterisk denotes significant ($p<0.05$) differences.

**Supplemental Figure 3 | Neural information in single units and power modulation by area**

A. Cue information, quantified with Percent Explained Variance (PEV) by area during the cue processing window (0.05 s-1s post cue onset). Mean +/- SEM across units. Red asterisk and horizontal black bars indicate significant differences in total information between each area and V4. B. Same as A, but for the pre-sample window (1.5 s – 0.05 s pre-cue onset). C-E, Percent change LFP power modulation by area during the sample window, separately by theta, alpha/beta, and gamma bands. Red asterisks indicate significant differences in power modulation between each area and V4.

**Supplemental Figure 4 | Unpredicted vs. predicted Power by area during the pre-sample window and area V4 selectivity in pre-sample window**

A-E, Percent change in power during the pre-sample window, comparing unpredicted vs. predicted power. Red bars denote unpredicted > predicted power, blue bars unpredicted < predicted power, $p<0.05$, cluster-based randomization test corrected for multiple comparisons. F-K, same as Figure 4, but for the pre-sample period.

**Supplemental Figure 5 | Unpredicted vs. predicted Coherence during the sample window**

Unpredicted minus predicted coherence, divided by the standard error of the mean for each area. Red bars denote unpredicted > predicted coherence, blue bars unpredicted < predicted coherence, $p<0.05$, cluster-based randomization test. Mean +/- SEM across all electrode combination for each inter-areal combination.

**Supplemental Figure 6 | Unpredicted vs. predicted Granger causality**

Same as Supplemental Figure 5, but for Granger causality, separately for both directions of Granger causal influence. The upper right interactions are feedforward, the lower left interactions are feedback.

### Supplemental Results

#### Relationship between spiking and LFP power

To test whether neurons with enhanced spiking to unpredictable samples tended to be at recording sites that also showed modulation of LFP power, we computed the correlation between modulation of spiking and LFP power by unpredictable vs unpredictable cuing from -1.25 to +1.25 seconds relative to cue onset. We took care to avoid selecting the same recording sites for spiking as for LFP power (to avoid spike-to-LFP bleedthrough, which would create spurious correlations). Instead, we compared a particular site's spiking to the LFP of the bipolar difference between LFPs at two adjacent recording sites (bipolar spacing was 400 um, see Methods). The correlation was performed for each electrode separately, and then averaged across areas.

In all areas, modulations of spiking and LFP gamma were positively correlated across sites (Supplemental Figure 2A). In the three areas with laminar-resolved recordings, the correlation between spiking and gamma modulation was stronger in superficial than deep layers (Wilcoxon rank sum test comparing unit-by-unit correlation between power and MUA in superficial vs. deep layers, $p<0.05$, Supplemental Figure 2B). In the alpha/beta-band, the correlation was negative in all areas (Supplemental Figure 2C) and stronger in deep cortical layers for areas V4 and 7A (Wilcoxon rank sum test comparing unit-by-unit correlation between power and MUA in superficial vs. deep layers, $P<0.05$, Supplemental Figure 2G). In the theta-band, effects by areas and layer were mixed. The correlation between spiking and LFP modulation was positive in V4 and not layer specific. It was around zero in 7A, and slightly negative in PFC superficial layers (Supplemental Figure 2D), where the correlation was different between superficial and deeper layers (Supplemental Figure 2D, Wilcoxon rank sum test comparing unit-by-unit correlation between power and MUA in superficial vs. deep layers, $P<0.05$).

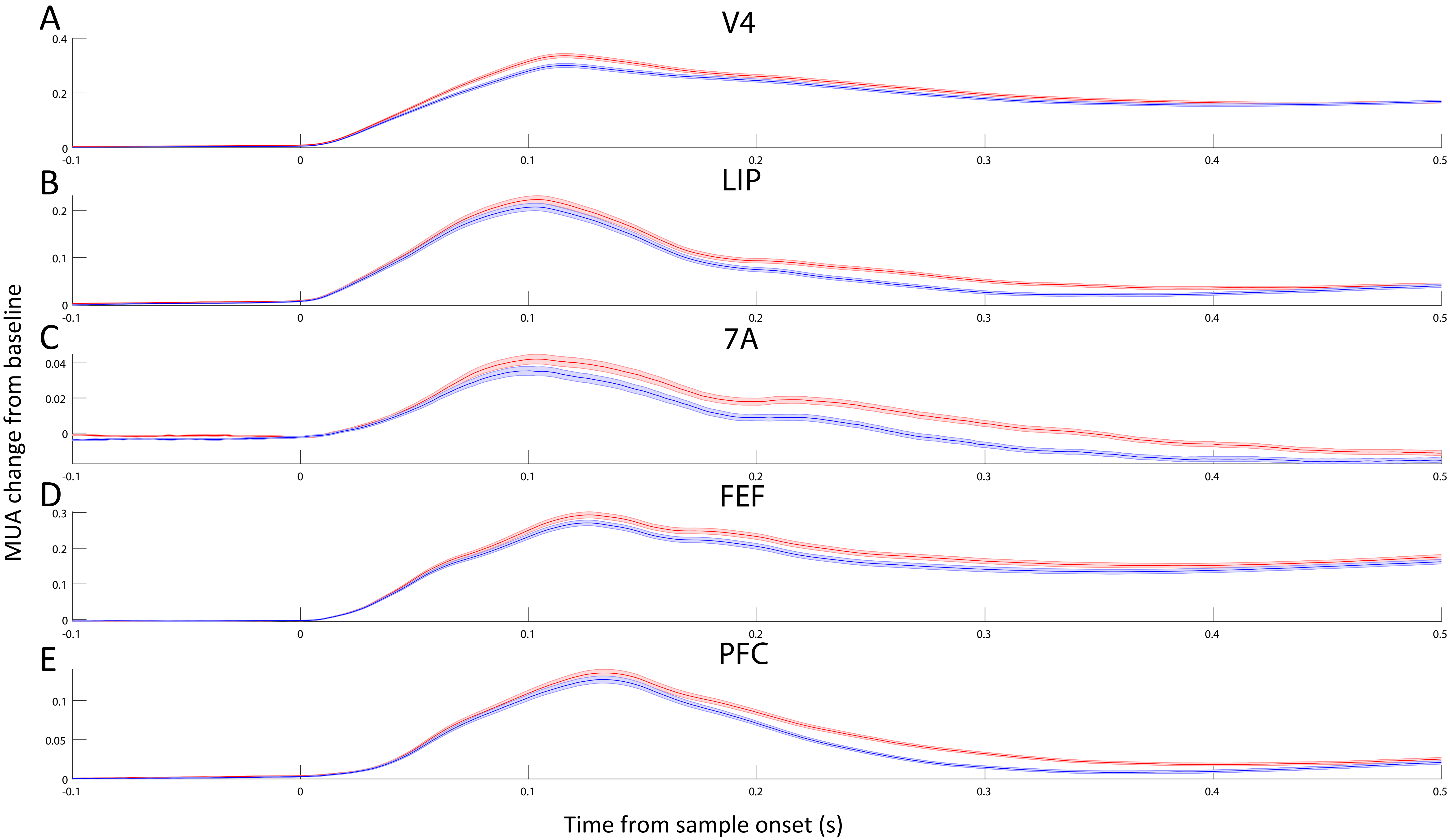
A
V4
B
LIP
C
7A
D
FEF
E
PFC
MUA change from baseline
Time from sample onset (s)
0.4
0.2
0.1
0.04
0.02
0.3
0.05
0
-0.1
0.5

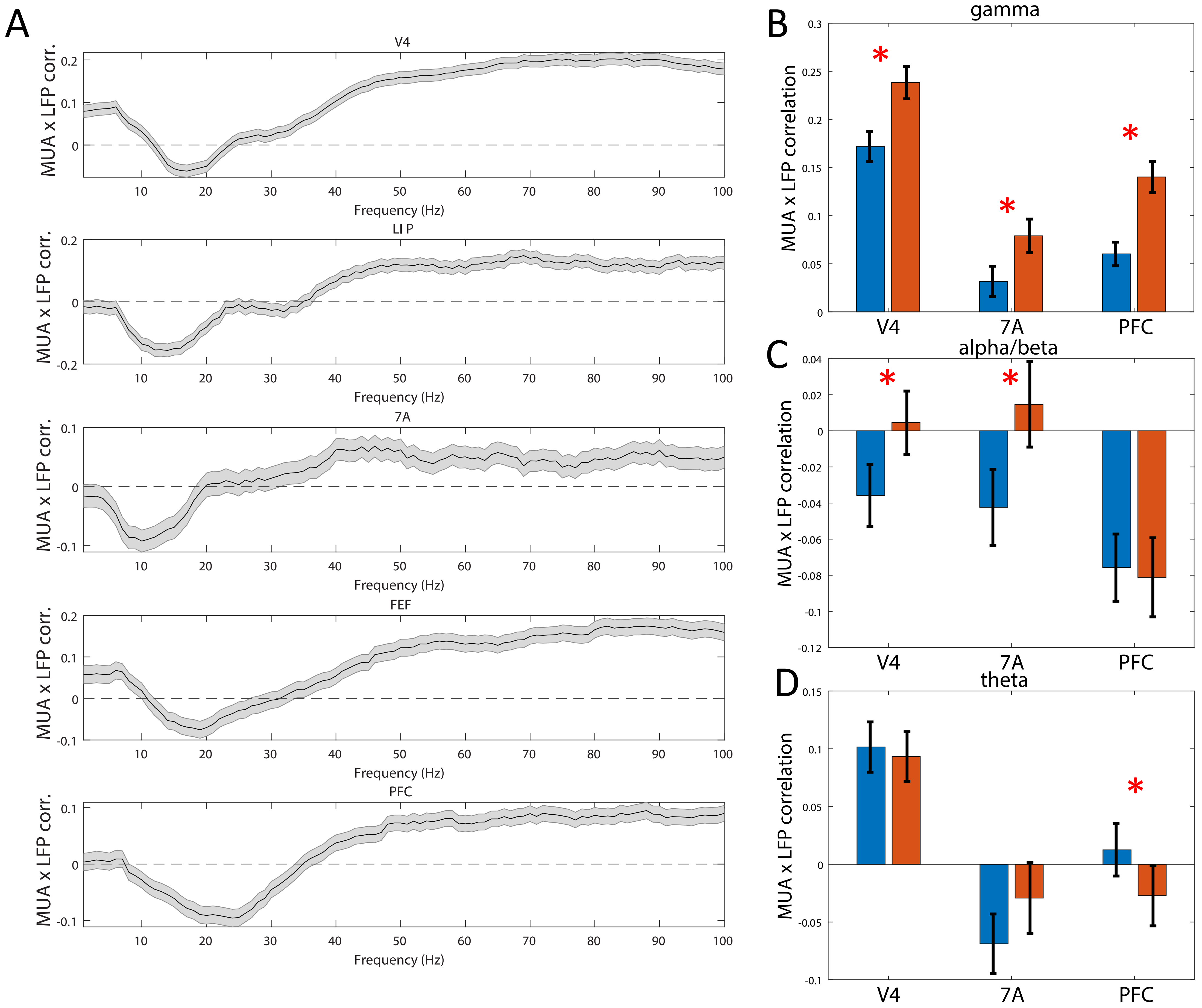

A
V4
LI P
7A
FEF
PFC
MUA x LFP corr.
Frequency (Hz)
B
gamma
C
alpha/beta
D
theta
MUA x LFP correlation
V4
7A
PFC

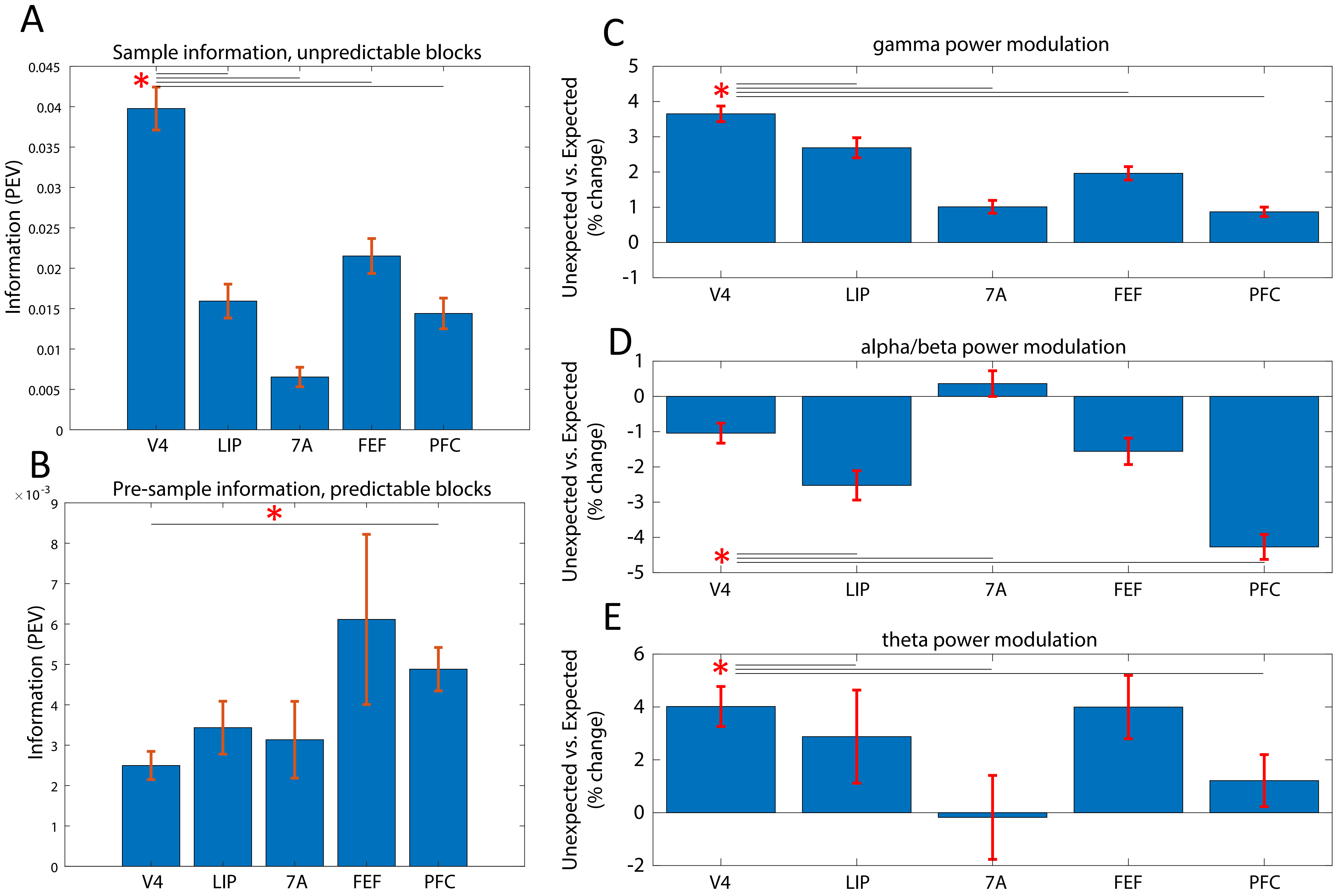

A
Sample information, unpredictable blocks
Information (PEV)
0.045
0.04
0.035
0.03
0.025
0.02
0.015
0.01
0.005
0
V4
LIP
7A
FEF
PFC
*
B
Pre-sample information, predictable blocks
×10^-3
Information (PEV)
9
8
7
6
5
4
3
2
1
0
*
C
gamma power modulation
Unexpected vs. Expected (% change)
5
4
3
2
1
0
-1
*
D
alpha/beta power modulation
1
0
-1
-2
-3
-4
-5
*
E
theta power modulation
6
4
2
0
-2
*

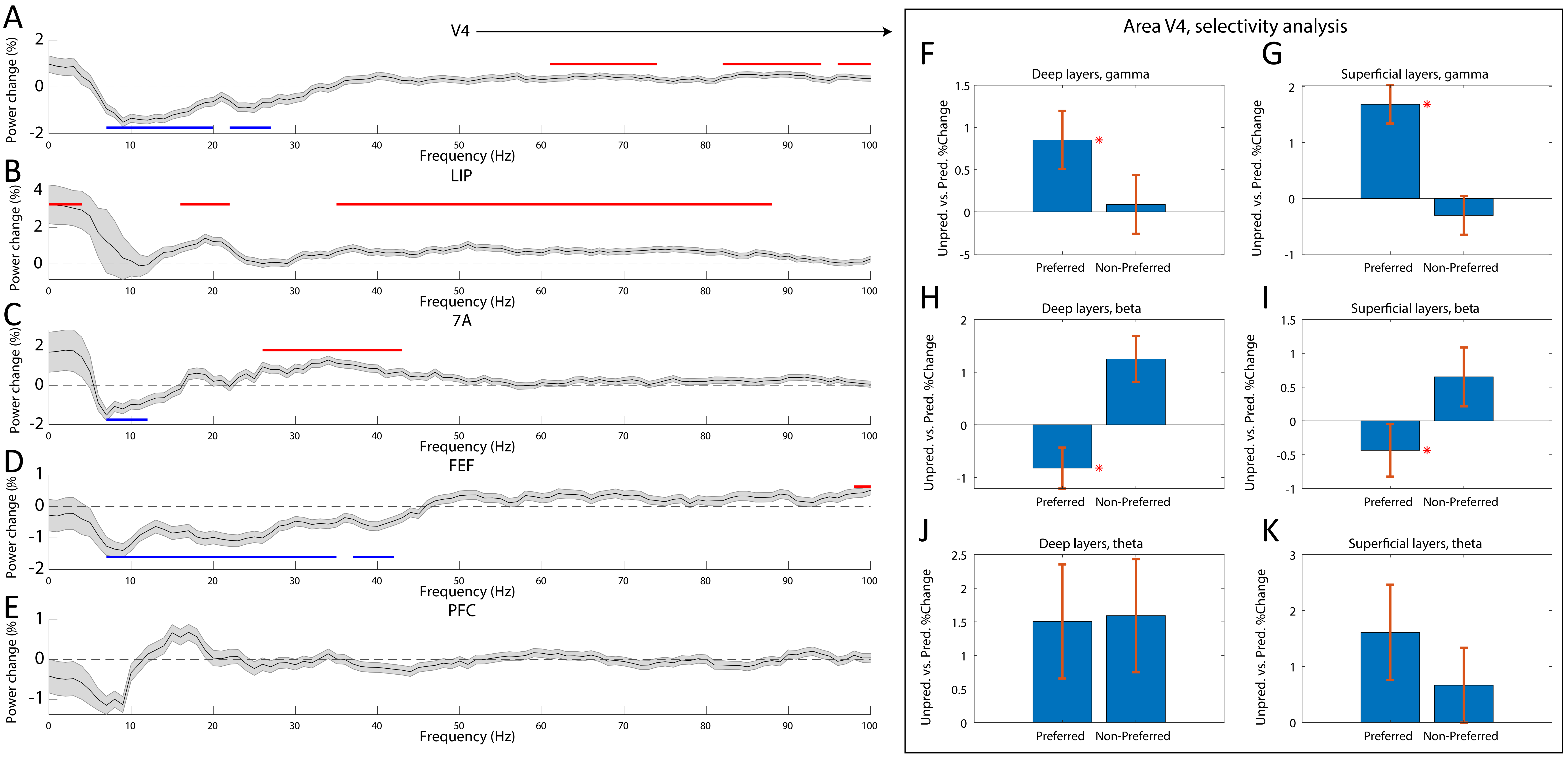

A
V4
Power change (%)
Frequency (Hz)
B
LIP
C
7A
D
FEF
E
PFC
Area V4, selectivity analysis
F
Deep layers, gamma
G
Superficial layers, gamma
H
Deep layers, beta
I
Superficial layers, beta
J
Deep layers, theta
K
Superficial layers, theta
Unpred. vs. Pred. %Change
Preferred
Non-Preferred

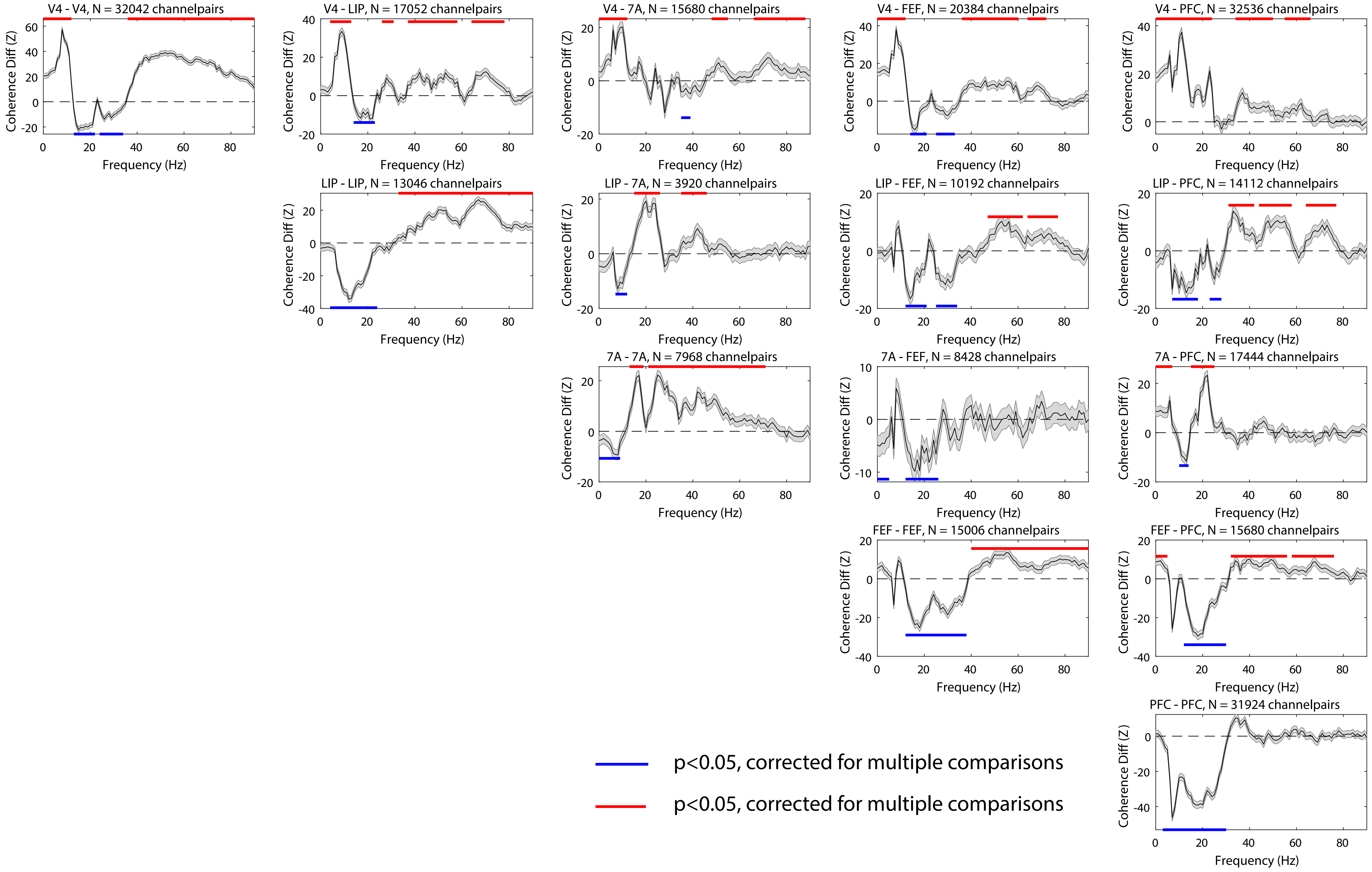

V4 - V4, N = 32042 channelpairs
V4 - LIP, N = 17052 channelpairs
V4 - 7A, N = 15680 channelpairs
V4 - FEF, N = 20384 channelpairs
V4 - PFC, N = 32536 channelpairs
LIP - LIP, N = 13046 channelpairs
LIP - 7A, N = 3920 channelpairs
LIP - FEF, N = 10192 channelpairs
LIP - PFC, N = 14112 channelpairs
7A - 7A, N = 7968 channelpairs
7A - FEF, N = 8428 channelpairs
7A - PFC, N = 17444 channelpairs
FEF - FEF, N = 15006 channelpairs
FEF - PFC, N = 15680 channelpairs
PFC - PFC, N = 31924 channelpairs
Coherence Diff (Z)
Frequency (Hz)
p<0.05, corrected for multiple comparisons
p<0.05, corrected for multiple comparisons

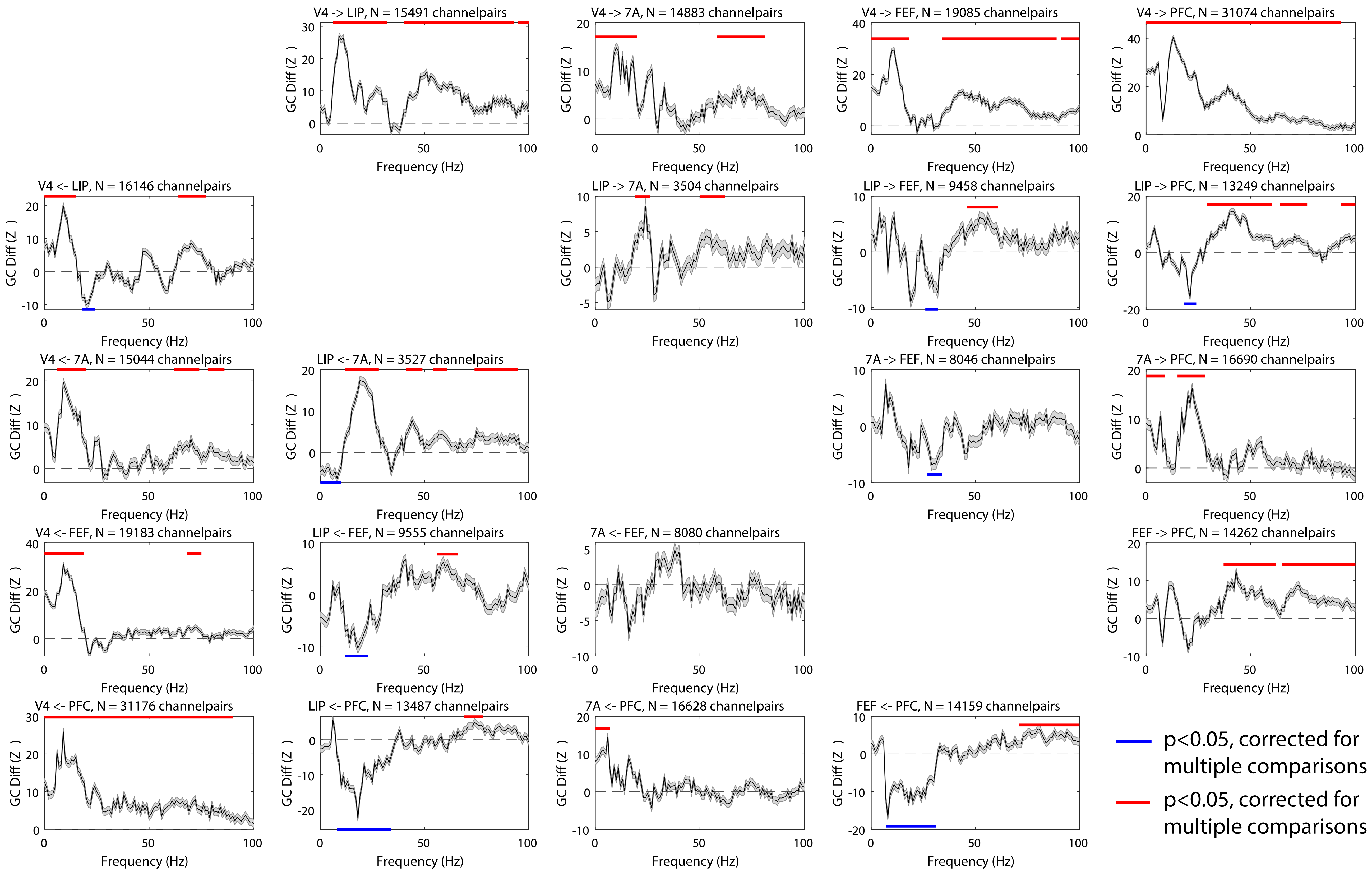

V4 -> LIP, N = 15491 channelpairs
V4 -> 7A, N = 14883 channelpairs
V4 -> FEF, N = 19085 channelpairs
V4 -> PFC, N = 31074 channelpairs
V4 <- LIP, N = 16146 channelpairs
LIP -> 7A, N = 3504 channelpairs
LIP -> FEF, N = 9458 channelpairs
LIP -> PFC, N = 13249 channelpairs
V4 <- 7A, N = 15044 channelpairs
LIP <- 7A, N = 3527 channelpairs
7A -> FEF, N = 8046 channelpairs
7A -> PFC, N = 16690 channelpairs
V4 <- FEF, N = 19183 channelpairs
LIP <- FEF, N = 9555 channelpairs
7A <- FEF, N = 8080 channelpairs
FEF -> PFC, N = 14262 channelpairs
V4 <- PFC, N = 31176 channelpairs
LIP <- PFC, N = 13487 channelpairs
7A <- PFC, N = 16628 channelpairs
FEF <- PFC, N = 14159 channelpairs
GC Diff (Z )
Frequency (Hz)
p<0.05, corrected for multiple comparisons
p<0.05, corrected for multiple comparisons
